# Heterosis of leaf and rhizosphere microbiomes in field-grown maize

**DOI:** 10.1101/2020.01.13.904979

**Authors:** Maggie R. Wagner, Joe H. Roberts, Peter Balint-Kurti, James B. Holland

## Abstract

- Macroorganisms’ genotypes shape their phenotypes, which in turn shape the habitat available to potential microbial symbionts. This influence of host genotype on microbiome composition has been demonstrated in many systems; however, most previous studies have either compared unrelated genotypes or delved into molecular mechanisms. As a result, it is currently unclear whether the heritability of host-associated microbiomes follows similar patterns to the heritability of other complex traits.
- We take a new approach to this question by comparing the microbiomes of diverse maize inbred lines and their F_1_ hybrid offspring, which we quantified in both rhizosphere and leaves of field-grown plants using 16S-v4 and ITS1 amplicon sequencing.
- We show that inbred lines and hybrids differ consistently in composition of bacterial and fungal rhizosphere communities, as well as leaf-associated fungal communities. A wide range of microbiome features display heterosis within individual crosses, consistent with patterns for non-microbial maize phenotypes. For leaf microbiomes, these results were supported by the observation that broad-sense heritability in hybrids was substantially higher than narrow-sense heritability.
- Our results support our hypothesis that at least some heterotic host traits affect microbiome composition in maize.

## INTRODUCTION

The role of host genotype in shaping microbiome composition and function is a crucial topic for both basic and applied research into plant-associated microbial communities. Understanding patterns and mechanisms of microbiome heritability is a key step toward understanding how plant-microbiome interactions evolved, and unlocking their potential to improve agricultural productivity and sustainability (Bordenstein & Theis, 2015; Busby *et al*., 2017). Heritable components of microbiome variation may reflect plant-microbe interactions that have been shaped by natural selection acting on plant traits underlying fitness, which might be modified during crop breeding. Plant genotype can also affect the phenotypic response to variation in the local microbial species pool. Thus, genetic variation within and among species may have major implications for harnessing the potential benefits of plant-associated microbiomes, which include improved nutrient acquisition, drought resistance, defense, and reproductive phenology (Friesen *et al*., 2011; Lau & Lennon, 2012; Wagner *et al*., 2014; Panke-Buisse *et al*., 2015, 2017; Santhanam *et al*., 2015; Busby *et al*., 2016; Ritpitakphong *et al*., 2016; Berg & Koskella, 2018; Hubbard *et al*., 2019).

Although many studies have documented host genetic variation for microbiome composition, most of these have compared genotypes whose relationship to each other is undefined (Bodenhausen *et al*., 2013; Edwards *et al*., 2015; Wagner *et al*., 2016; Wallace *et al*., 2018a; Walters *et al*., 2018) or delved into molecular mechanisms using mutants, transgenic manipulation, or QTL-mapping approaches (Bressan *et al*., 2009; Bodenhausen *et al*., 2014; Horton *et al*., 2014; Beckers *et al*., 2016; Ritpitakphong *et al*., 2016). As a result, we have limited ability to predict microbiome similarity between two host genotypes that are directly related *(e.g*., within a known pedigree). Recently, the introgression of quantitative trait loci was shown to alter leaf microbiomes across five generations of a breeding experiment for improved disease resistance in maize (Wagner *et al*., 2020); however, additional studies of this type are needed before we will be able to anticipate what microbiome changes might follow genetic improvement in crops or evolution in natural populations.

One major outstanding question is whether hybridization between genetically distinct lineages results in hybrids that harbor microbiomes that are intermediate in composition between those of the parent genotypes. This question is important because intermating between diverged populations or species can create novel combinations of alleles that modify adaptively important traits, with major evolutionary implications (Anderson & Stebbins, 1954; Soltis & Soltis, 2009). Hybridization is also the cornerstone of many crop breeding strategies. In many cases, and particularly in maize, traits of hybrids exceed both of their parents rather than being intermediate between their parents. This results in more robust, productive plants—a phenomenon known as heterosis or hybrid vigor.

Although heterosis has been the subject of intense research for over 100 years, its causes and mechanisms are still not completely understood. It likely involves various genetic mechanisms including dominance, overdominance, and genic dosage, and affects a wide variety of morphological and physiological traits (Birchler *et al*., 2003; Flint-Garcia *et al*., 2009; Schnable & Springer, 2013). To the extent that host traits affecting microbiome composition exhibit heterosis, microbiome composition itself could potentially do the same (Figure 1). So far, however, exploration of the links between heterosis and plant microbiomes has been limited to specific microbial groups and has excluded aboveground plant-microbe interactions. For instance, maize hybrids are more likely than inbreds to be colonized by arbuscular mycorrhizal fungi and by beneficial *Pseudomonas* strains that produce growth-promoting phytohormones and protective antibiotic compounds in the rhizosphere (Picard *et al*., 2004, 2008; Picard & Bosco, 2005, 2006; An *et al*., 2010).

**Figure 1.**
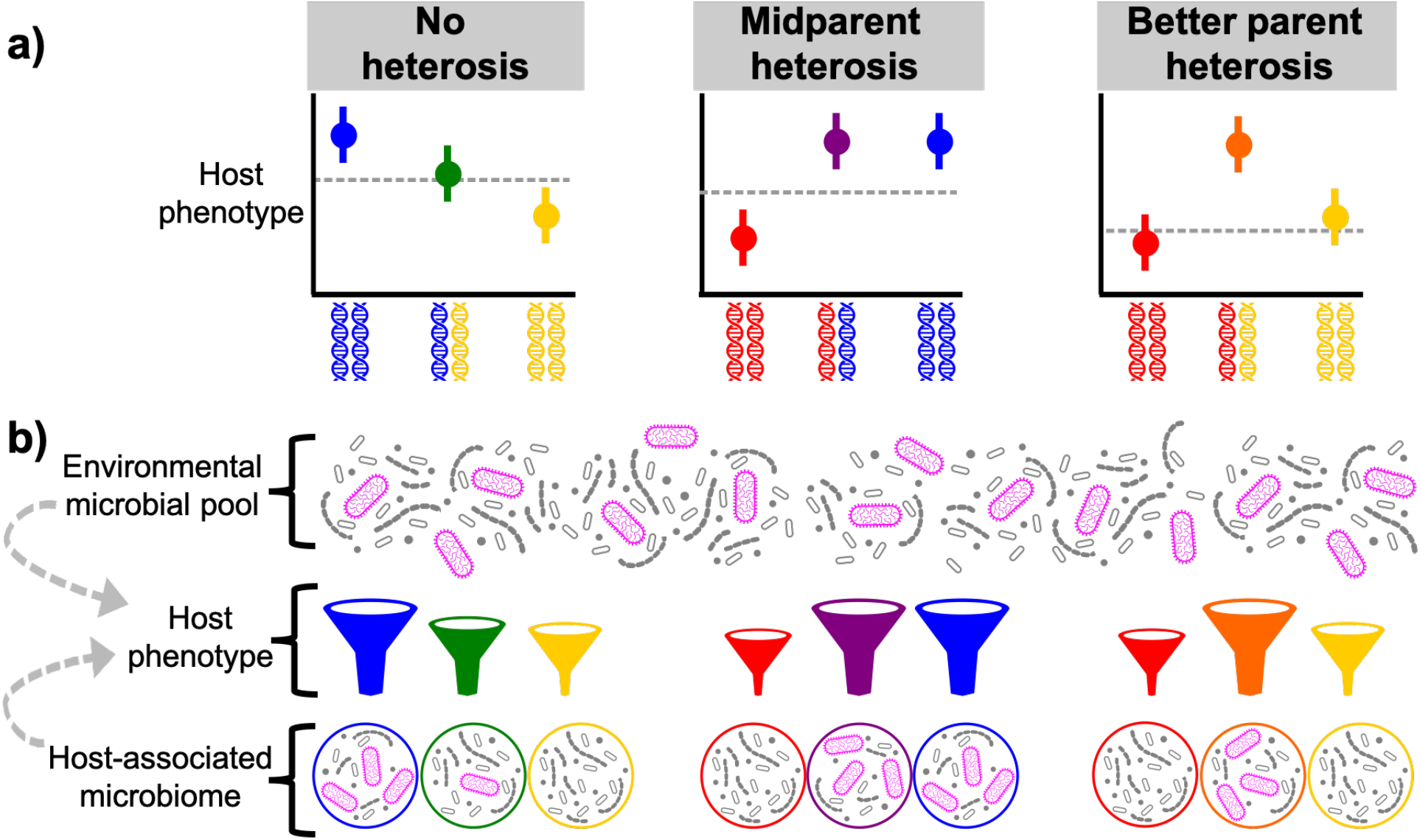
Overview of heterosis and its hypothesized relationship to microbiome composition. **(a)** Each panel depicts trait values for each of three genotypes: two inbred lines (left and right, symbolized by blue, yellow, or red in the different panels) and their F_1_ hybrid (center, symbolized by green, purple, or orange). The dotted grey lines denote the midparent value, or the expected trait value of the hybrid in the absence of heterosis. *Midparent heterosis* describes hybrid trait values that deviate from the midparent value but are within the range of the parent line values. *Better parent heterosis* describes hybrid trait values that are outside of the range of parental values. **(b)** Host phenotype defines the habitat available to potential microbial colonizers from the ambient community. Therefore, heterosis for traits that affect microbiome composition can result in heterosis for microbiome composition itself (*e.g*., relative abundance of the focal microbe highlighted in pink). The dashed grey arrows represent the complication that microbes can alter phenotypic expression in the host.

Here, we investigate the effects of hybridization on maize leaf and rhizosphere microbiomes. In maize, variation for leaf and rhizosphere microbiome composition has been identified among diverse inbred lines (Peiffer *et al*., 2013; Wallace *et al*., 2018a; Walters *et al*., 2018). The genes and traits underlying this variation are unknown, and hybrid offspring of these or other inbred lines have not been fully characterized for microbiome content. Here, we focus on linking host genotype to the microbiome, treating the phenotypic mechanisms as a “black box”; this experiment was not optimized to identify the plant traits affecting microbial symbionts. We used amplicon sequencing to quantify bacterial and fungal communities in eleven F_1_ hybrids and their parent lines, grown together in two experimental fields. We address the following questions: (1) Do the microbiomes of maize hybrids differ consistently from those of inbred lines? (2) Which microbiome features, if any, show heterosis within individual crosses? and (3) Are patterns of microbiome heritability in F_1_ crosses comparable to those of other complex traits? The results are a significant step towards understanding the relationship between host genotype and microbiome composition.

## MATERIALS AND METHODS

### Field experimental design

In May 2017, 18 maize genotypes were planted in a randomized design into two fields at Central Crops Research Station, Clayton, NC. The fields were 1.1 km apart and had similar soil types but different recent crop rotation histories and soil microbiota (Tables S1–S2).

The genotypes included the inbred lines B73 and Mo17; five additional inbred lines selected from the founding members of the Nested Association Mapping population (Yu *et al*., 2008); and 11 F_1_ hybrids resulting from crosses of those lines to B73 and Mo17 (Figure 2a). For all hybrids, either B73 or Mo17 was the maternal parent; for the B73xMo17 hybrid, B73 was the maternal parent. We included B73 and Mo17 due to their importance as elite representatives of the stiff-stalk and non-stiff-stalk heterotic groups and because they have high-quality reference genomes that will facilitate deeper dissection of host genetic microbiome control (Jiao *et al*., 2017; Sun *et al*., 2018). The five additional inbred lines were chosen because they showed highly divergent rhizosphere microbiome composition relative to each other and to B73 and Mo17 in a previous study (Peiffer *et al*., 2013). Some of these were sweetcorn lines or tropical in origin; these groups are distinct from B73 and Mo17 in a wide variety of phenotypic traits including root system architecture and phenology, all of which could potentially influence microbiome composition. Thus, our study captured a wide range of maize genetic and phenotypic variation; however, identifying the particular traits that caused any observed microbiome changes was beyond the scope of this experiment.

**Figure 2.**
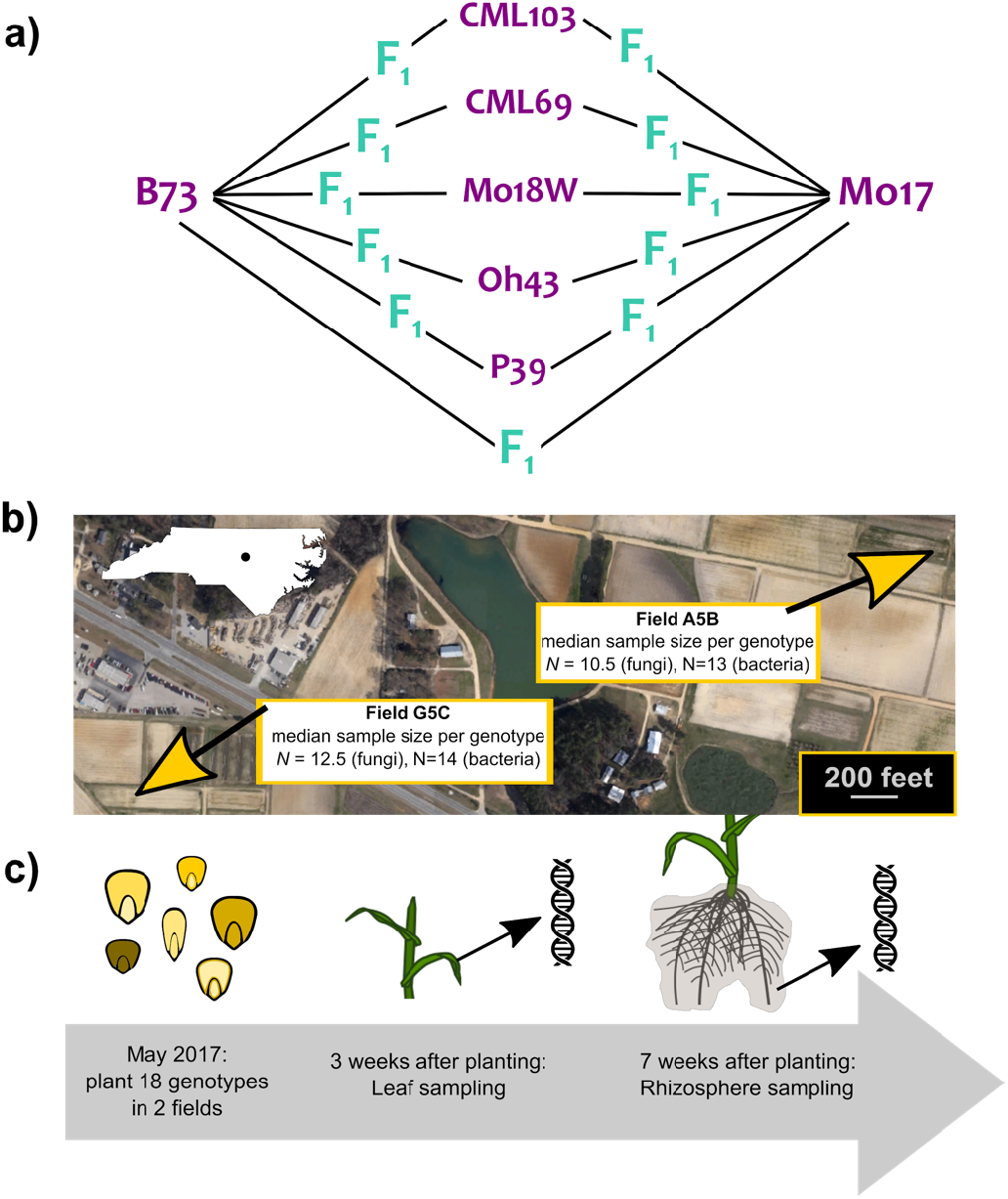
Overview of experimental design. **(a)** Seven inbred lines and eleven F_1_ hybrids (with B73 or Mo17 as the common maternal parent) were represented in our experiment. **(b)** Twenty to thirty replicates of each genotype were planted in randomized blocks in each of two fields at Central Crops Research Station, Clayton, NC, USA. Due to uneven germination and survival, the median sample size per genotype was slightly lower in Field A5B than in Field G5C; due to uneven sequencing of 16S and ITS amplicons and subsequent quality filtering, the median sample size was slightly lower for the fungal dataset than the bacterial dataset. Map data: Google (2019). **(c)** Leaf discs were collected from young plants three weeks after planting. Plants were uprooted seven weeks after planting (shortly before anthesis for most genotypes) and rhizosphere was collected and pooled from all root types to a depth of ~1 foot.

Immediately prior to planting, kernels were soaked in 3% hydrogen peroxide for two minutes to reduce surface-associated microbes and then rinsed in distilled water. One seed per genotype was planted into each block, except for B73 and Mo17 which were planted an average of 1.5 times per block. Twenty blocks of 18 or 20 individuals, randomized with respect to genotype, were planted into adjacent rows within each field. However, the final sample sizes were lower due to uneven germination, survival, and sequencing (Figure 2b).

### Sample collection

Three weeks after planting, we sampled leaf tissue from all emerged plants for microbiome analysis. We chose this early timepoint to ensure that sampling was completed before the establishment of foliar diseases such as southern leaf blight, which sometimes arise later in the season in these fields and which could potentially skew microbiome composition (but see (Wagner *et al*., 2020)). A standard hole punch and forceps (rinsed in 70% ethanol between samples) were used to collect three discs evenly spaced from base to tip of the fourth leaf, avoiding the midrib and any non-green tissue. Leaf discs were immediately placed into sterile tubes and kept on ice until transfer to −20°C for storage.

Seven weeks after planting, we uprooted plants for collection of rhizosphere soil. We chose this timepoint so that our data would be comparable to the largest published maize rhizosphere dataset available at the time (Peiffer *et al*., 2013). This timepoint was originally chosen based on the expectation that host genetic effects would accumulate in the rhizosphere microbiome until the transition to flowering, while the roots were mostly a carbon sink (Peiffer & Ley, 2013). Using a spade, we collected root crowns (~6” deep x ~18” wide) and shook them until little or no bulk soil was falling from the roots. We then used ethanol-rinsed pruners to cut all roots starting 2” below the top of the crown. All whorls of roots attached to the stem up to 6” belowground were included in each sample; for large root systems that could not be grasped in one hand, we instead cut all roots from one hemisphere of the crown to ensure that primary, seminal, and crown roots would be represented in each sample. A variety of root system properties likely differed among genotypes from different heterotic groups, or between hybrids and inbreds; however, we conducted this experiment agnostic to the particular phenotypic traits that affect microbiome composition. We therefore consider such genetic variation in root morphology or physiology to be a possible underlying cause of any observed microbiome variation, which would need to be verified through additional experimentation.

Cut roots were stored in a sterile plastic bag on ice until transfer to −20°C for storage. Rhizosphere collections were spread over four days (July 2^nd^-5^th^, 2017), with representatives of all genotypes harvested from each field on each day so that collection date would not be confounded with genotype or field (Figure S1). At time of rhizosphere sampling most plants had transitioned to adult phase but were not yet flowering, with the exception of the sweet corn P39 which began anthesis during the collection period. We also collected bulk soil samples from the four corners and approximate center of each field, approximately 8 cm below the surface. The first ten soil samples were collected on July 4^th^, and sampling was repeated on July 5^th^ for a total of 20 samples.

We vortexed roots in 50 mL of 1x phosphate-buffered saline (PBS) at maximum speed for 15 s to dislodge rhizosphere soil. The resulting suspension was poured through a sterile 100-micron stainless steel mesh into a sterile 50 mL tube, then centrifuged for 30 minutes at 1,600 x *g* to concentrate the rhizosphere soil. We then discarded the supernatant, resuspended rhizosphere pellets in 1.5 mL 1x PBS, transferred them to sterile Eppendorf tubes, and centrifuged them for 5 minutes at 10,000 x *g*. After discarding the remaining supernatant we transferred rhizosphere pellets into storage at −80°C until DNA extraction.

To determine whether this procedure affected downstream data collection and inference, we tested the protocol on our bulk soil samples. Each bulk soil sample was divided into two aliquots. We extracted DNA directly from one aliquot; to the other, we first applied the same washing protocol that we used to separate rhizosphere from roots. Comparison of data from washed and unwashed bulk soil samples indicates that the procedure may have resulted in underestimates of alpha diversity and overestimates of beta diversity (i.e., resulted in loss of some taxa and generated additional variation among samples) (Figure S2). Alternatively, alpha diversity in bulk soils may have been inflated by the presence of dead cells or loose DNA that were separated out by the washing process. Our data on rhizosphere microbiomes should be interpreted with these caveats in mind.

### DNA extraction

Prior to DNA extraction, we vortexed leaf discs at top speed for 10 seconds in 750 μL sterile deionized water to remove dust and soil. Leaf discs were shaken dry and blotted on clean paper towels, frozen at −80°C, and lyophilized. Freeze-dried leaf discs were randomly arranged within 96-well plates and ground for 60 s at 25 Hz in a Retsch MM301 mixer mill. As a positive control we included a mock microbial community in three wells (ZymoBiomics Microbial Community Standard, Zymo Research, Irvine, CA, USA). We used the Synergy 2.0 Plant DNA Extraction Kit (OPS Diagnostics, Lebanon, NJ, USA) to purify DNA, doubling the manufacturer’s recommended bead-beating time to improve lysis. Purified DNA was eluted in 30 μL of 1x TE buffer (pH 8.0).

We used the MoBio PowerSoil htp-96 kit to extract DNA from rhizosphere and bulk soil samples. Rhizosphere pellets were resuspended in 750 μL bead solution and transferred to 96-well plates, with samples randomly arranged across plates. To improve lysis, samples were subjected to two freeze-thaw cycles prior to extraction. Washed and unwashed bulk soil samples (see above) were randomly distributed among the rhizosphere samples.

### Amplicon library preparation and sequencing

As in previous work (Wagner *et al*., 2020), we characterized bacterial and fungal communities by sequencing the V4 region of the 16S rDNA gene and the ITS1 region of the fungal rDNA gene, respectively. We amplified these barcoding genes using the primer pairs 515f/806r and ITS1f/ITS2, modified to include 3 to 6 random nucleotides upstream of the gene primers to increase library complexity (Lundberg et al. 2013). A second PCR step added dual-indexed Illumina adaptors to the amplicons. Barcoding genes were amplified in 10 μL reactions that included 5 μL of 5Prime HotMasterMix (Quanta Bio, Beverly, MA, USA) and 0.4 μL of each 10 μM primer. For 16S-V4 amplification from leaf samples, we added 0.15 μL of PNA (100 μM) to block amplification of host mitochondrial and chloroplast sequence (Lundberg *et al*., 2013).

The template for the first PCR step was 1.5 μL of DNA extracted from leaf, rhizosphere, or mock community samples; for the second PCR step, the template was 1 μL of the products from the first step. Reaction conditions for ITS1 amplification were 2 minutes at 95°C; 27 cycles of 20 s at 95°C, 20 s at 50°C, and 50 s at 72°C; and a final 10 minutes at 72°C. Conditions for 16S-V4 amplification were identical except for an additional PNA-annealing step (5 s at 78°C) before primer annealing at 52°C. The first PCR step included 27 cycles while the second included 8 cycles. PCR products from the first step were purified using 7 μL of magnetic SPRI bead solution, two washes with 70% ethanol, and elution in 10 μL water. Finally, we pooled 1 μL of each dual-indexed library, creating four separate pools (i.e., fungal and bacterial libraries for leaf and rhizosphere samples). We used Blue Pippin (Sage Science Inc., Beverly, MA, USA) to select fragments that were between 150 bp and 600 bp, and then combined the four pools at equimolar concentrations. The final combined sample was sequenced on the Illumina HiSeq 2500 platform (250-bp paired-end reads).

### Sequence processing and quality filtering

We used CUTADAPT v1.12 (Martin, 2011) to trim primers from raw sequences prior to quality filtering. Using DADA2 v1.10.1 (Callahan *et al*., 2016), we removed all reads with ambiguous bases or more than 2 expected errors. Forward and reverse 16S reads were truncated at 220 bp and 160 bp (respectively). We inferred error rates separately for 16S and ITS data using 3×10^6^ bases, then de-replicated and de-noised reads to identify amplicon sequence variants (ASVs). We removed chimeric ASVs using DADA2 and then used the RDP classifier (Wang *et al*., 2007) trained on the RDP training set (v. 16) and the UNITE database (supplemented with *Zea* sequences) to assign taxonomy to bacterial and fungal ASVs, respectively (Cole *et al*., 2014; Nilsson *et al*., 2019).

We discarded “non-usable” ASVs that could not be reliably identified at the kingdom level and those identified as host sequences. Samples with <500 usable reads were excluded from analysis. Within each sample we removed bacterial and fungal ASVs with less than 0.21% and 0.185% relative abundance, respectively. These cutoffs were determined using the mock community positive controls to identify relative abundance thresholds that minimized cross-contaminant ASVs while retaining true positives and discarding as little data as possible. We required ASVs to be observed at least 25 times in at least 5 samples to be considered “reproducible”; all others were removed from the dataset (Lundberg *et al*., 2012). Combined, these quality filters reduced the number of bacterial ASVs from 8,958 to 753 and the number of fungal ASVs from 4,863 to 501. However, 92.3% of the original bacterial data and 95.0% of the fungal data were retained. The final number of observations for each sample was normalized, centered, and used to control for sampling effort in downstream analyses.

### Quantification of fungal abundance in the rhizosphere

For rhizosphere and soil samples, we used droplet digital PCR (ddPCR) to measure the number of fungal 18S rRNA genes per unit of extracted DNA. We interpret this metric as an estimate of the absolute abundance of fungi (*cf*. bacteria and other organisms) within the community. DNA extracts were quantified using the PicoGreen dsDNA Assay Kit (Invitrogen, Carlsbad, CA, USA) according to the manufacturer’s instructions, and all samples were diluted to a working concentration of 0.102 ng/uL. We used the FungiQuant assay to quantitatively amplify the fungal 18S rRNA gene (Liu *et al*., 2012). Each ddPCR reaction mixture contained 1 ng template DNA (9.8 uL of the diluted working solution), 5 uL of 2x Evagreen Probe Supermix (Bio-Rad, Hercules, CA, USA), 0.1 uL of each 100nM primer (from 5’ to 3’, forward primer GGRAAACTCACCAGGTCCAG and reverse primer GSWCTATCCCCAKCACGA), and molecular-grade water to bring the reaction volume to 20uL. The reaction mixtures were loaded into DG8 droplet generation cartridges along with 70uL of droplet generation oil for EvaGreen assays, and the QX200 Droplet Generator was used to divide the mixture into droplets (Bio-Rad). The droplets were then transferred to a T100 thermal cycler (Bio-Rad) for 10 minutes of activation at 95°C; followed by 50 cycles of denaturation (30s at 95°C), annealing (60s at 55°C), and elongation (30s at 72°C); and finally held at 4C until use. After amplification, the droplets were transferred to the QX200 Droplet Reader and the QuantaSoft AP program (Bio-Rad) was used to calculate the number of fungal 18S rRNA gene copies per ng of rhizosphere DNA.

### Data analysis

Data analysis was performed in parallel for bacteria and fungi using R version 3.6.0 with the packages phyloseq, vegan, DESeq2, tidyr, lme4, and lmerTest (McMurdie & Holmes, 2013; Love *et al*., 2014; Bates *et al*., 2015; Kuznetsova *et al*., 2017; Wickham, 2017; Oksanen *et al*., 2019). Leaf and rhizosphere communities were analyzed in parallel. Raw sequence reads are available at the NCBI Sequence Read Archive under Bioproject #PRJNA597058. Processed data, metadata, and original code are available in a Zenodo archive (doi: 10.5281/zenodo.3596465).

ASV counts were normalized using the variance-stabilizing transformation from the DESeq2 package (Love *et al*., 2014). Before normalization we estimated alpha (within-sample) diversity using the abundance-based coverage estimator (ACE) and Shannon metrics (Hughes *et al*., 2001). We quantified beta (between-sample) diversity by calculating the centroid for each genotype in each field in multivariate space; then, using the function “betadisper” from the package vegan (Oksanen *et al*., 2019), we determined the distance of each sample to its respective centroid.

#### Comparing microbiome features of hybrid vs. inbred maize

First, we investigated whether the microbiomes of maize hybrids are distinct from those of inbreds. We modeled alpha diversity (ACE and Shannon metrics) and beta diversity (distance to centroid) using separate linear mixed models, with “category” (*i.e*., either hybrid or inbred), field, and their interaction as fixed effects. Normalized sequencing depth was included as a covariate. Genotype (nested within category), block (nested within field), and plate (to control for batch effects) were included as random-intercept terms. For rhizosphere samples, collection date was also included as a random effect. ANOVA with Type III sums-of-squares and Satterthwaite’s approximation of degrees-of-freedom was used for statistical inference of fixed effects; likelihood ratio tests were used for random effects.

To test whether hybrids and inbreds differed in overall community composition, we conducted permutational MANOVA of Bray-Curtis dissimilarities, with hybrid/inbred category, field, and their interaction as predictors. Sequencing depth and collection date were included as nuisance variables. Finally, we fit negative binomial models to explore which microbial taxa differed in relative abundance between hybrids and inbreds. ASV counts were modeled with hybrid/inbred category as the predictor (Love *et al*., 2014; McMurdie & Holmes, 2014). The rarest taxa were excluded from this analysis: the threshold for inclusion was set at 10% of the mean abundance. For example, the mean ASV count across the leaf fungal dataset was 14,057, so we included only ASVs that were observed at least 1,406 times. This reduced the number of tests by 45% but still used 98.3% of the data. *P*-values from Wald tests were adjusted to correct for multiple comparisons using the false discovery rate (Benjamini & Hochberg, 1995).

#### Testing for microbiome heterosis

Next, we explored patterns of heterosis within individual crosses for various microbiome features, including alpha and beta diversity (as described above); overall community composition (summarized using the top 5 unconstrained principal coordinates and the top 5 constrained axes of variation from a distance-based redundancy analysis constrained on genotype); and variance-stabilized counts of ASVs. For each feature we fit a linear mixed-effects model with the predictors field, sampling effort, batch effects, and (for rhizosphere samples) collection date. The purpose of these models was not to test for significance, but to derive residual trait values cleansed of these sources of noise; the resulting residuals were then used to test for evidence of heterosis.

For each feature, we calculated the mean values of these residuals in the seven inbred lines, from which we calculated the expected mid-parent values (assuming additive genetic variance) for each hybrid. We then conducted two-sided *t*-tests of the null hypothesis that each hybrid’s microbiome trait value was equivalent to its respective mid-parent value (i.e., tests for “mid-parent heterosis”; Figure 1a). In addition, we tested for “better-parent heterosis” using one-sided *t*-tests to assess whether the hybrid value fell outside of the parental range. Finally, we used linear regression to test for a relationship between microbiome heterosis (percent deviation from expected mid-parent value) and average heterozygosity, estimated using genome-wide SNP data from the maize HapMap v2 (Chia *et al*., 2012).

#### Effects of host relatedness on microbiome similarity within and between environments

To test whether the relationship between two host genotypes predicts the similarity of their microbiomes, we calculated the Bray-Curtis dissimilarity between all pairwise combinations of samples. Each pair of samples was classified according to the known relationship (or lack thereof) between the two host genotypes. We fit a linear model of Bray-Curtis dissimilarity with host-pair relationship as the predictor. The log-transformed absolute difference in sequencing depth between samples was included as a covariate; host genotypes and individual plant IDs were included as random intercept terms to account for these sources of non-independence.

We used a similar approach to assess whether the microbiomes of hybrids and inbred lines differed in their sensitivity to environmental variation. We modeled Bray-Curtis dissimilarity between pairs of individuals with the same genotype using hybrid/inbred category, within/between fields, and their interaction as predictor variables. The log-transformed absolute difference in sequencing depth between samples was included as a covariate; the host genotype and the two individual plant IDs were included as random intercept terms.

#### Estimating heritability, general combining ability (GCA), and specific combining ability (SCA) for microbiome composition

Finally, we used ASReml-R software to estimate GCA and SCA for key properties of hybrid microbiomes, including alpha diversity, beta diversity, and the five major constrained and unconstrained axes of variation from the dbRDA described above (Isik *et al*., 2017). To estimate variance components for each of these microbiome properties, we fit separate linear mixed-effects models for inbreds and hybrids with field, sequencing depth, and collection date as fixed effects. For the inbred model, the random effects included batch, block nested within field, genotype, and genotype-field interactions. For the hybrid model, random effects included batch, block, female parent, male parent, the interaction of each parent line with field, the interaction between the parent lines, and the three-way interaction between the parent lines and field.

For hybrids, a diallel model was fit to the data, in which a single GCA effect was estimated for each parent line, regardless of whether it was a male or female parent (Möhring *et al*., 2011). GCA variance is related to the variation of parental breeding values in hybrids and additive variance was estimated as twice the GCA variance. SCA variance was calculated as the variance component due to the interaction between the parent lines and represents variation of dominance genetic effects, such as those that could lead to heterosis. The total genetic variation was calculated as the sum of twice the GCA plus SCA; the total genotype-by-environment (GxE) variation was calculated as the sum of the variance components from interactions between each parent line and field (twice the GCA-field interaction plus the SCA-field interaction variances). Narrow-sense heritability (*h*^2^) was calculated as the additive variance divided by the total phenotypic variation; broad-sense heritability (*H*) was calculated as the total genetic variation (2*GCA + SCA) divided by the phenotypic variation. For inbreds, total genetic variation and GxE were calculated using the variance components for genotype and genotype-by-field interaction, respectively.

## RESULTS

The final dataset included 501 fungal ASVs and 753 bacterial ASVs. The fungal dataset comprised 481 leaf samples and 332 rhizosphere samples from a total of 547 individual plants, with a median sampling effort of 19,266 observations per sample. The bacterial dataset included 502 leaf samples and 403 rhizosphere samples from 567 individuals (median sampling effort = 9,055 observations per sample). Plant survival differed between fields and genotypes, but the median sample size ranged from *N* = 10.5 to *N* = 14 per genotype per field (Figure 2).

Leaf and rhizosphere communities differed substantially in diversity and composition (Figure S3, Figure S4). Leaf communities were dominated by *Pantoaea, Entyloma*, and *Golubevia* spp., while the most abundant taxa in the rhizosphere were *Ralstonia, Burkholderia, Rhizobium, Entyloma*, and *Golubevia* (Table S3). Only 193 bacterial ASVs were observed in both sample types; 260 bacterial ASVs were found only in leaves, whereas 290 were specific to the rhizosphere. For fungi, 132 ASVs were specific to leaves, 196 were found only in the rhizosphere, and 173 were found in both (Figure S4). Alpha diversity was higher in the rhizosphere than in leaves. For fungi, beta diversity was also higher in the rhizosphere, suggesting that belowground fungal communities may be more strongly influenced by microhabitat variation or stochastic processes (Figure S3c-d). This is consistent with the observation that community composition differed strongly between fields only for rhizosphere fungi (Figure S3a-b). As expected, rhizosphere communities were compositionally distinct from bulk soil (Figure S2); they also were relatively richer in fungi (Figure S5).

### Microbiome composition differs between maize hybrids and inbred lines

First, we investigated whether hybrids, as a group, differed from inbreds in microbiome diversity and composition. For bacteria and leaf-associated fungi, we found no consistent difference between these groups in either alpha diversity or beta diversity (Table S4). In contrast, fungal communities in the rhizospheres of hybrid plants had higher alpha diversity (within-sample species richness and evenness) and lower beta diversity (among-individual variability) relative to inbreds (Tukey’s HSD, *P* < 0.05).

Hybrids’ microbiomes were primarily distinguished from those of inbred plants not by alpha or beta diversity, but by overall community composition (Table 1). Distance-based redundancy analysis (ordination of Bray-Curtis dissimilarity, constrained to emphasize variation explained by hybrid/inbred category) revealed that microbiomes of maize hybrids clustered with those of other hybrids, apart from those of inbreds (Figure 3). We confirmed this observation using permutational MANOVA; hybrid/inbred category predicted the composition of leaf-associated fungal communities and rhizosphere bacterial and fungal communities (ADONIS, all *P*<0.01). This differentiation was strongest in the rhizosphere, explaining about 1.6% of the overall variation (compared to <1% in leaves). Using negative binomial models, we identified 18 bacterial and fungal ASVs that were differentially abundant in the rhizosphere of hybrid versus inbred hosts (Figure 4).

**Figure 3.**
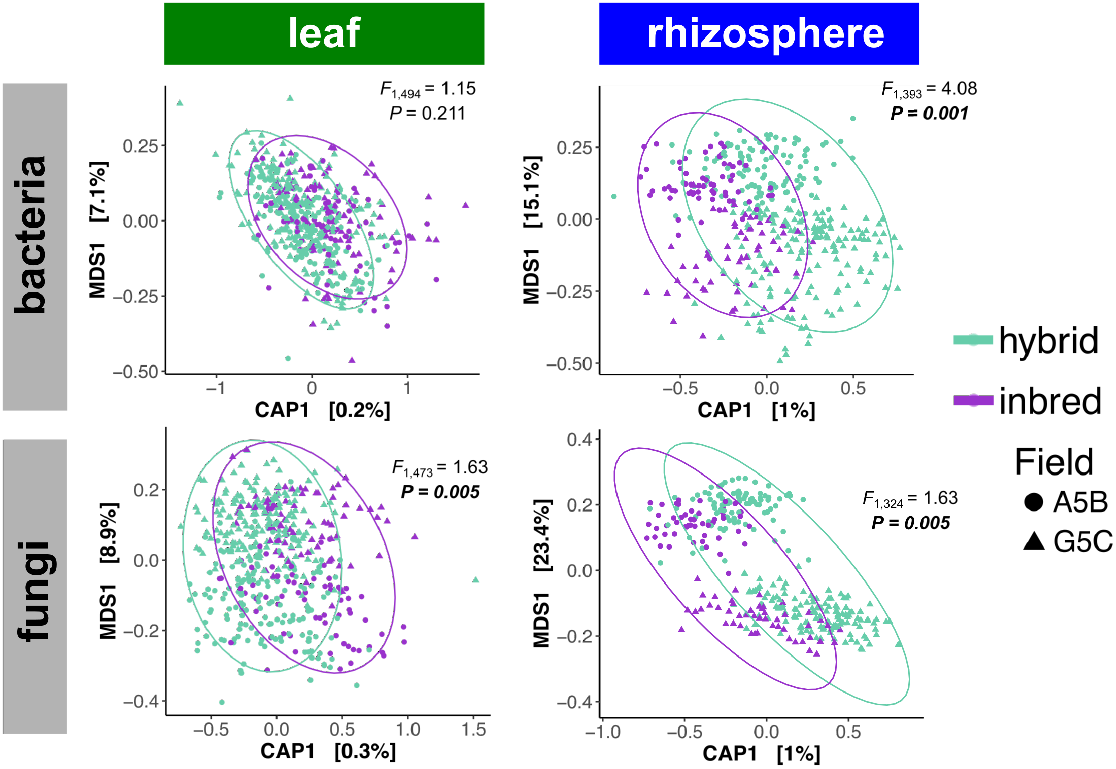
Constrained distancebased redundancy analysis revealed that the microbiome composition of maize hybrids is distinct from that of inbred lines. Two axes of variation in community composition (ordination of Bray-Curtis dissimilarity) are plotted for bacterial and fungal communities in maize leaves and rhizosphere. The CAP1 axis was constrained to capture variation explained by inbred/hybrid category; MDS1 was the dominant unconstrained axis of variation. ANOVA-like permutation tests were performed to assess statistical significance of the constrained axis.

**Figure 4.**
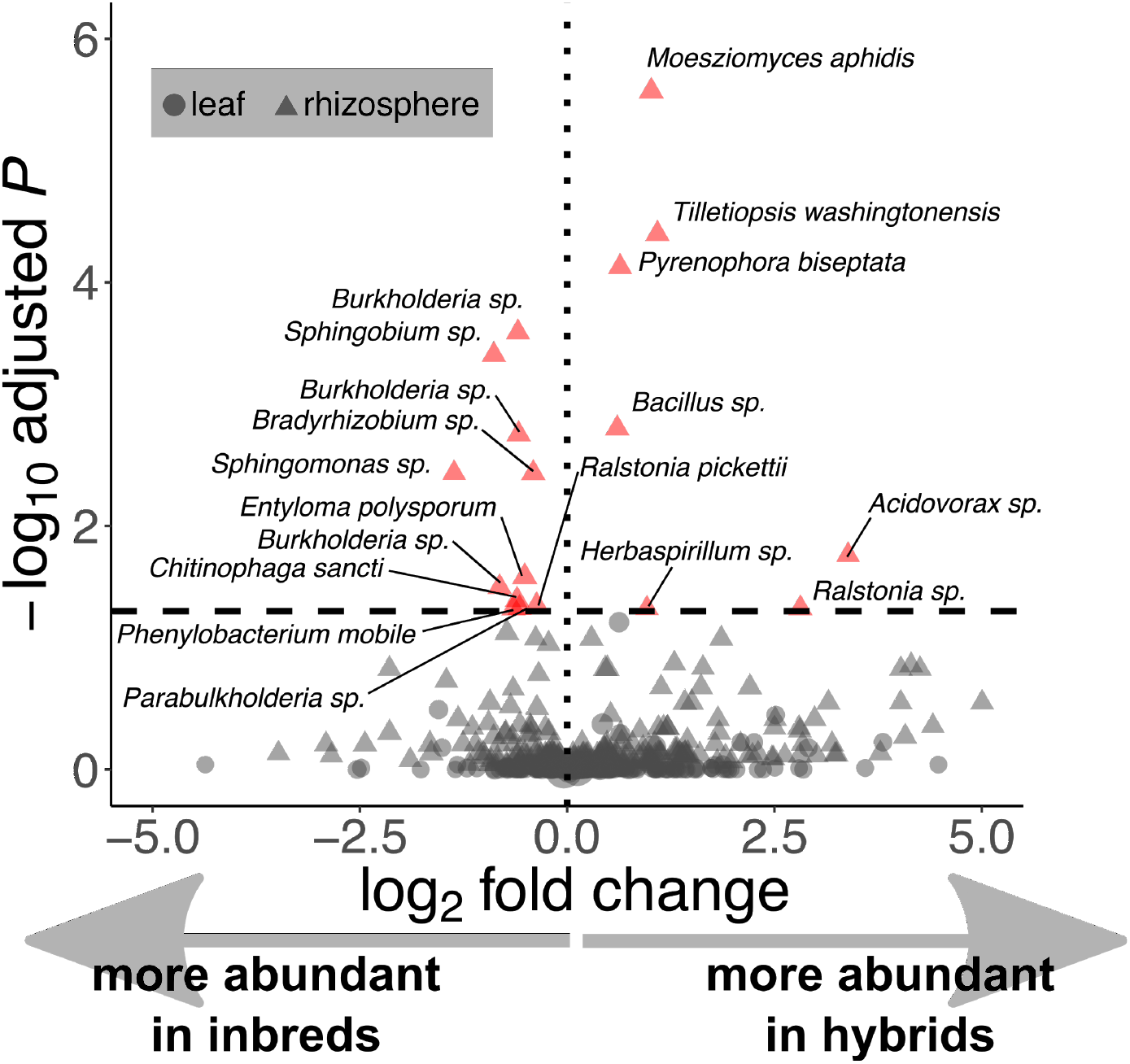
Diverse bacterial and fungal taxa were enriched or depleted in the rhizosphere of maize hybrids relative to inbred lines. For this analysis, data from the two fields were pooled. Each point represents one ASV, with point size scaled by the relative abundance of that taxon. FDR-corrected *P*-values from Wald tests of negative binomial models of taxon counts are plotted against the estimated effect sizes from the same models. Red points reflect taxa that were significantly associated with host hybrid/inbred category (FDR < 0.05). ASVs are labeled as the closest-matching species based on comparison to the NCBI sequence database using BLAST. One leaf ASV (*Microdochium sorghi*) is omitted to improve figure clarity; it was strongly enriched in hybrids.

**Table 1.**
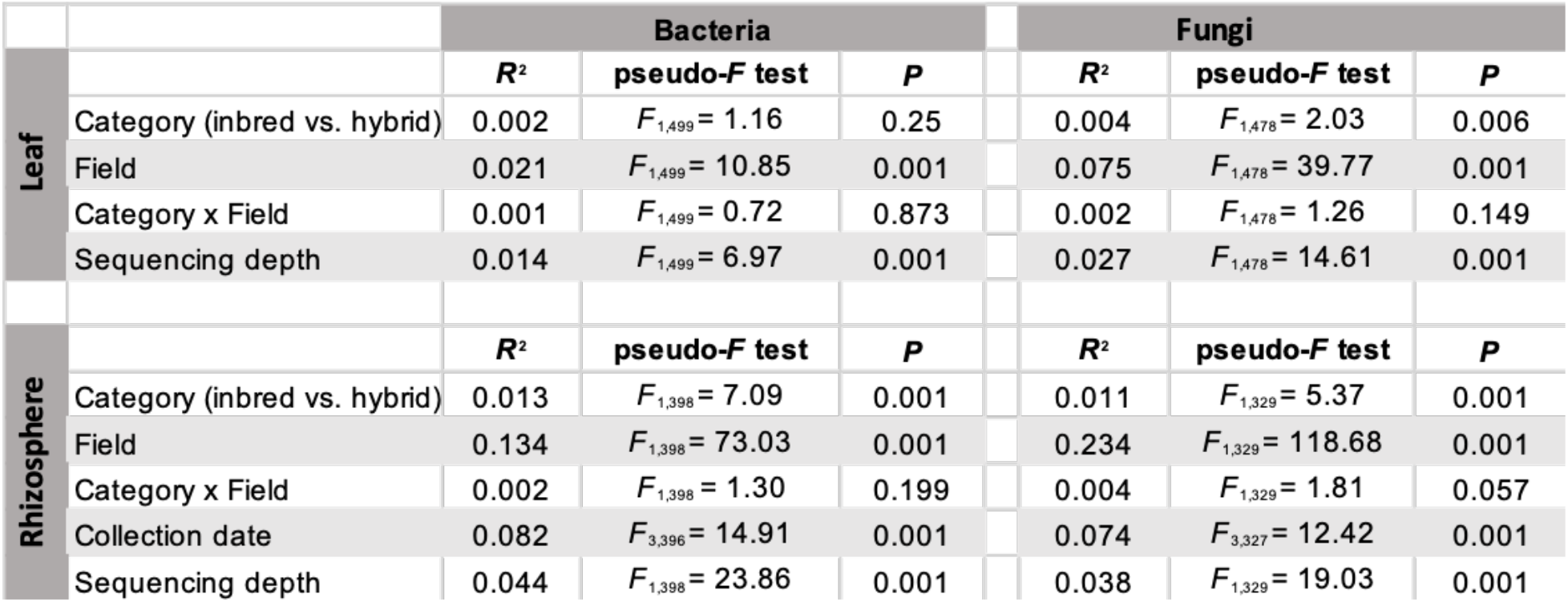
Permutational MANOVA of fungal and bacterial community composition in maize leaves and rhizosphere. *P*-values were generated from 999 permutations of Bray-Curtis dissimilarities among samples, which were calculated based on the matrix of variance-stabilized counts of amplicon sequence variants (ASVs). Results of an analogous test for variation among genotypes within hybrid/inbred categories are provided in Table S7.

### Patterns of heterosis in microbiome features

Next, we tested which microbiome features, if any, showed evidence of heterosis within individual crosses. These features included community-level metrics of alpha and beta diversity (*i.e*., Shannon, ACE, distance-to-centroid) and composition (the top 5 constrained and unconstrained axes of variation from distance-based redundancy analysis; Figure 5b). We used *t*-tests to assess whether these microbiome features deviated from the midparent expectation for each hybrid, using a significance threshold of FDR < 0.05 (Figure 1). We note that historically, the term “better-parent” heterosis describes cases in which the hybrid has higher values of a desirable trait or lower values of an undesirable trait relative to both parents (Flint-Garcia *et al*., 2009). Because the function and value of these microbiome traits (if any) are unknown, we use this term to refer to cases in which hybrid trait values are higher than both parents or lower than both parents (Figure 1a).

**Figure 5.**
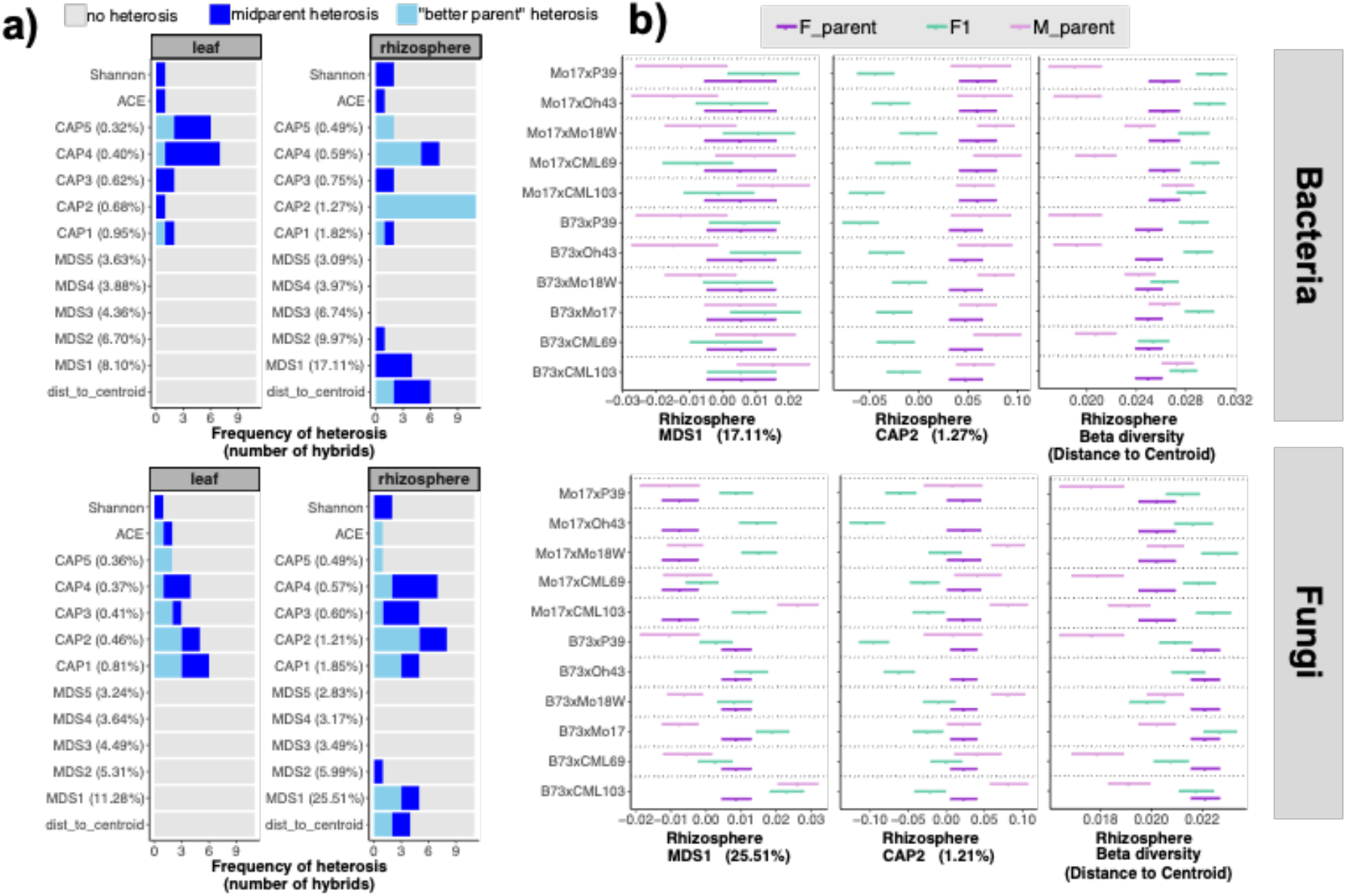
Multiple microbiome properties show evidence of heterosis in maize leaves and rhizosphere for both bacteria (top of each panel) and fungi (bottom of each panel). We calculated the values of select microbiome features in maize hybrids, after controlling for field, sequencing depth, and batch effects; then, we used *t*-tests to compare hybrid trait values to the mid-parental trait values as illustrated in Figure 1. *P*-values were adjusted to correct for multiple tests using the false discovery rate (Benjamini & Hochberg, 1995). **(a)** Heterosis was more common for some microbiome properties than others. This difference is especially stark for community composition, summarized using the top 5 constrained and unconstrained axes of variation resulting from distance-based redundancy analysis. The constrained axes (abbreviated “CAP” for Constrained Analysis of Principal Coordinates) capture the microbiome variation that is associated with host genotype; the unconstrained axes (abbreviated “MDS” for Multidimensional Scaling) capture the dominant axes of remaining variation. The percentage of the overall community composition explained by each CAP or MDS axis is specified on the *y*-axis. **(b)** Patterns of heterosis are illustrated for two axes of variation in rhizosphere community composition, as described for panel (a), and for rhizosphere beta diversity (distance to centroid), which quantifies within-group/among-individual variation in community composition. Points represent least-squares mean values +/- 1 s.e.m.

Heterosis for alpha and beta diversity was detected in a few crosses, but the most frequently heterotic microbiome features were the axes of variation in community composition. Both midparent and “better-parent” heterosis were particularly common in the dbRDA axes that were constrained by host genotype (Figure 5a), which is consistent with these axes capturing the heritable components of microbiome composition. More microbiome features displayed heterosis in the rhizosphere than in leaves, which is consistent with the overall stronger differences between hybrids and inbreds observed previously (Figure 3). Linear regressions revealed that genome-wide heterozygosity did not reliably predict the strength of microbiome heterosis, quantified as the percent deviation from expected midparent values of microbiome features (Figure S6). Heterozygosity was positively associated with heterosis only for the bacterial MDS1 in the rhizosphere (*R*^2^ = 0.53, *P* = 0.03). For bacterial CAP1 and fungal beta diversity, the relationship was inverse. Most frequently, however, we found no significant relationship, although this could reflect low statistical power due to the small number of hybrids tested (*N* = 11).

We also tested for heterosis of individual ASVs using FDR-corrected *t*-tests of their variance-stabilized abundances. In most hybrids, most ASVs did not deviate from the midparent expectation (Figure S7a). However, 40% of rhizosphere fungal ASVs showed evidence of heterosis in at least one cross, as did 20% of leaf-associated fungal ASVs, representing a wide diversity of classes (Figure S7b; Table S5). For bacterial ASVs, the rates of heterosis in the rhizosphere and leaf were 35% and 17%, respectively. The prevalence of ASV heterosis varied among hybrids, ranging from 27% of the tested fungal ASVs in the B73xP39 rhizosphere, to 5.1% of the tested bacterial ASVs in the B73xMo17 rhizosphere (Figure S7a). The most strongly heterotic fungal ASV (showing “better-parent” heterosis most frequently) belonged to the yeast *Moesziomyces aphidis*, which was enriched in the rhizospheres of six hybrids relative to both of their inbred lines (Figure S8). Two bacterial ASVs in the rhizosphere (*Pseudonocardia* sp. and *Sphingomonas* sp.) and one in the phyllosphere (*Acinetobacter* sp.) each showed “better-parent” heterosis in five hybrids (Figure S8).

For both bacteria and fungi, the second constrained axis of rhizosphere variation (CAP2) was the most frequently heterotic microbiome property; hybrid values of bacterial CAP2 exceeded both parental values for all 11 crosses (Figure 5). To obtain a biological interpretation of CAP2, we identified the top 10% of ASVs with the strongest loadings on this axis (Table S6). These included 13 of the ASVs that were differentially abundant between hybrids and inbred lines (Figure 4), including three *Burkholderia* ASVs. Additional rhizosphere ASVs with strong influences on this axis included three more *Burkholderia* ASVs, two *Nitrospira* ASVs, and four ASVs from the family Rhizobiaceae.

### Patterns of microbiome heritability in F_1_ crosses

Finally, we investigated general patterns of microbiome heritability in this set of maize crosses. In particular, we (1) partitioned variance in community composition among genotypes within hybrid/inbred categories; (2) compared microbiome similarity of pairs of genotypes with differing levels of relatedness; (3) tested whether the microbiomes of hybrid and inbred maize were equally influenced by environmental variation; and (4) investigated the relative importance of additive versus heterotic components of genetic variation for shaping hybrid microbiomes.

First, we used permutational MANOVA to test for host genetic variation among inbred lines and among hybrids for whole-community composition. Host genotype contributed 4-8% of the variation in overall community composition for both bacteria and fungi, in both leaves and rhizosphere (Table S7). Alpha and beta diversity also varied among host genotypes, but not as consistently as community composition. For instance, host genotype affected alpha diversity and distance to centroid (*i.e*., among-individual variation within genotypes); but almost exclusively in the rhizosphere (Table S4).

Next, to assess how relatedness affects microbiome similarity, we modeled Bray-Curtis dissimilarity between pairs of samples as a response to host-genotype-relationship using ANOVA. Overall, host-genotype-relationship had a stronger effect on rhizosphere microbiome dissimilarity than leaf microbiome dissimilarity (Figure 6a). Bacterial rhizosphere composition differed more between pairs of hybrids (regardless of their relatedness) than between pairs of inbreds; the opposite was true in leaves. We found no evidence that hybrids’ microbiomes are more similar to those of their maternal inbred than to their paternal inbred; in fact, for leaf-associated bacteria, the opposite was true. Similarly, hybrid microbiomes were equally dissimilar to those of their half-siblings regardless of whether the female or male parent was shared (Figure 6a).

**Figure 6.**
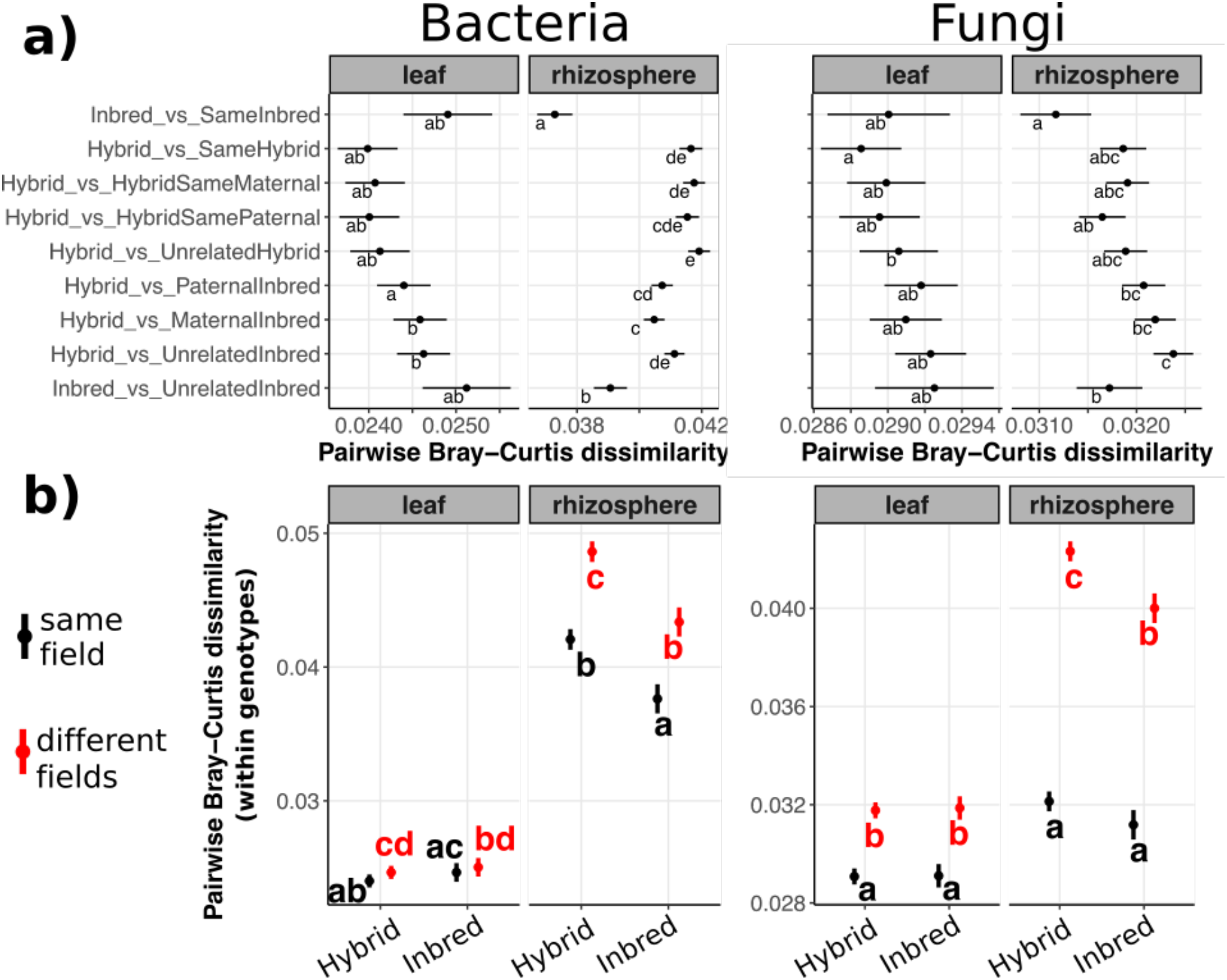
Host relatedness affects dissimilarity between pairs of microbiomes. **(a)** Bray-Curtis dissimilarity (horizontal axis) was calculated for each pair of samples, each of which was categorized according to the relationship between the two samples’ host genotypes (vertical axis). All sample pairs were from individuals in the same field. We used ANOVA to test the response of Bray-Curtis dissimilarity to “host-pair-relationship”; the log absolute value of the difference in sequencing depth between the samples was included as a nuisance variable. Least-squares mean dissimilarity values from this model are shown +/- 1 s.e.m. Non-overlapping sets of letters indicate post-hoc Tukey’s Honest Significant Differences. **(b)** Pairs of rhizosphere samples from inbred genotypes were more similar to each other in composition than pairs of samples from hybrid genotypes. Bray-Curtis dissimilarity was modeled as a function of “inbred/hybrid category”, “within vs. between field”, and their interaction; least-squares means are plotted as described in (a).

Third, we tested whether genotype-by-environment (GxE) interactions are more prominent for inbred microbiomes than hybrid microbiomes, as they are for many plant traits (Lewis, 1954; Griffing & Zsiros, 1971; Blum, 2013; Zanewich *et al*., 2018) (but see (Li *et al*., 2018)). We found that pairs of siblings from hybrid genotypes had *less* similar rhizosphere microbiomes than pairs of siblings from inbred genotypes, both within fields and between fields (Figure 6b). In addition, fungal load in the rhizosphere was stable across fields for inbreds, but not hybrids (Figure S5). For leaf microbiomes, however, there was no difference in sibling-pair dissimilarity between inbreds and hybrids. The variance components contributed by Genotype and GxE supported these findings. For many leaf microbiome traits, G/GxE was equal between hybrids and inbreds; for others (particularly fungal traits) it was higher in hybrids, and for others (particularly bacterial traits) the opposite was true (Figure S9a).

Last, we investigated whether hybrid microbiomes were influenced more by general combining ability (GCA; the average trait values of the two parent lines, mostly involving additive gene effects) or specific combining ability (SCA; the interaction between the two parent lines, involving synergistic or heterotic effects). Using this method we detected moderate heritability for a variety of leaf microbiome features, but no heritability for rhizosphere microbiomes. For both fungi and bacteria, narrow-sense heritability (*h*^2^) was much higher among inbred lines than among hybrids (Figure S9b). For hybrids, comparison of SCA and GCA revealed that for multiple axes of leaf microbiome variation, broad-sense heritability was considerably higher than *h*^2^ (Figure S9c). This indicates that leaf microbiomes, in particular, were shaped by specific combinations of parental genotypes (*i.e*., heterotic or synergistic effects) beyond the general combining ability (additive effects) of those parents.

## DISCUSSION

Understanding patterns and mechanisms of microbiome heritability is critical to understanding the evolution of host-microbiome symbiosis in both natural and human-directed systems. Here we demonstrate that microbiome diversity and composition display some of the same patterns as other complex plant traits within F1 crosses. In general, leaf and rhizosphere microbiomes of hybrid maize differ reliably from those of inbred maize lines, much like performance-related traits such as height and yield (Flint-Garcia *et al*., 2009). Within crosses, we detected evidence for heterosis of a variety of microbiome features, including beta diversity and several orthogonal axes of community composition (Figure 5).

Overall, effects of hybridization (and of host genotype more generally) were stronger in the rhizosphere than in the leaves. The reasons for this are unclear. One possible explanation is that the difference in sampling time (3 weeks after planting for leaves, 7 weeks for rhizosphere) allowed more time for deterministic effects of host genotype to shape rhizosphere communities, whereas the leaf microbiome reflected more of the stochastic variation that contributes to initial community assembly (Maignien *et al*., 2014). It is important to note that in our study, the observed differences between leaf and rhizosphere communities reflect not only the distinction between aboveground and belowground organs, but also differences in abiotic environment and in plant developmental stage that correspond to the two timepoints. Maize leaf microbiomes change drastically over the growing season (Wagner *et al*., 2020). Nevertheless, studies from diverse plant species confirm that leaf and rhizosphere microbiomes are reliably distinct (Coleman-Derr *et al*., 2016; Wagner *et al*., 2016; Wallace *et al*., 2018b). In maize, individual leaves can senesce within weeks, especially early in development; in contrast, much of the root system remains active throughout the plant’s lifespan (Fusseder, 1987). Thus, maize leaves are generally more transient habitats for microbes than maize rhizosphere. However, the same is true for the perennial forb *Boechera stricta*, in which host genotype effects were stronger in leaves than in roots (Wagner *et al*., 2016). One potential explanation for this opposite result is that maize forms partnerships with mycorrhizal fungi, while (like other mustards) *B. stricta* does not; this hints at fundamental differences in mechanisms of interaction with belowground microbes.

In this study, host genotype and its interaction with environment explained between 4-8% of the overall variation in microbiome composition (Table S3). This is a high proportion relative to other studies of host genetic microbiome control, but low relative to studies of other maize traits; flowering time and resistance to various diseases often have heritabilities above 80% (Robertson *et al*., 2006; Buckler *et al*., 2009). However, this may be an underestimate of the true importance of host genotype. If only a subset of the microbiome is actually responsive to host variation, then using the entire community as a response variable is likely diluting the host genotype signal. In support of this possibility, some studies, including in maize, have found that heritability of microbiome functional potential *(i.e*., metagenome content) is higher than the heritability of taxonomy-based microbiome features such as those used in our analysis (Lemanceau *et al*., 2017; Wallace *et al*., 2018a).

Maize is well-known for the strength of heterosis in intraspecific crosses, and our results indicate that this phenomenon can be seen within leaf and rhizosphere microbiomes. However, not all maize crosses are equally heterotic, either for yield or for the underlying traits (Flint-Garcia *et al*., 2009; Guo *et al*., 2019). This, too, was evident in our microbiome data (Figure 5; Figure S7), although we found no consistent relationship between heterozygosity (estimated from genome-wide SNP data) and microbiome heterosis (Figure S6). However, with only 11 crosses included in our study, an explicit test for a relationship between the extent of microbiome heterosis and the extent of heterosis for other phenotypic traits was beyond the scope of this work. One hurdle to addressing this question is the general lack of knowledge about which plant traits cause changes in the microbiome. Mutant and transgenic studies in Arabidopsis have demonstrated that properties of the leaf cuticle affect microbial colonization (Bodenhausen *et al*., 2014; Ritpitakphong *et al*., 2016), and that physiological traits such as hormone signalling and defensive secondary chemistry influence the root and rhizosphere microbiomes (Bressan *et al*., 2009; Lebeis *et al*., 2015). However, it is unknown whether the same traits are involved in shaping the maize microbiome. An additional complication is the fact that the microbiome can, in turn, affect phenotypic expression in the host plant (O’Brien *et al*.; Panke-Buisse *et al*., 2017; Berg & Koskella, 2018) as well as the relationship between phenotype and yield (Wagner *et al*., 2014; Chaney & Baucom, 2020). For these reasons, careful experimentation—not just simultaneous measurement of plant traits and microbiome features— will be needed to clarify the phenotypic links between heterozygosity and microbiome heterosis (Figure 1).

Although no comparable studies are currently available, we expect that other sets of maize genotypes, and indeed other plant species, could show different effects of hybridization on the microbiome. A comparison to plant species that display outbreeding depression rather than heterosis would be particularly interesting. Evidence from animal systems suggests a potential role for the microbiome in speciation. Crosses between two subspecies of house mice (*Mus musculus*) result in hybrid offspring with reduced fertility and gut microbiomes that diverge from both parent subspecies (Wang *et al*., 2015), while lethality of hybrids between species of *Nasonia* wasps can be attributed to a disordered microbiome (Brucker & Bordenstein, 2013). To our knowledge, however, our study is the first to investigate whole microbiome composition in true intraspecific crosses of any organism. More data is needed from similar experiments, particularly in additional plant species that vary in mating system.

Notably, the lines used in our study are known to vary in phenology and developmental trajectories (Buckler *et al*., 2009). To the extent that the phenotypic traits affecting the microbiome vary over development, this variation may be one of the mechanisms underlying the observed microbiome differences among genotypes. This is more likely to be the case for rhizosphere samples (collected ~7 weeks after planting) than for the leaf samples (collected ~3 weeks after planting), because genetic variation for phyllochron is quite low in maize (Van Esbroeck *et al*., 2008). A weekly time-series revealed genotype-by-timepoint interactions influencing rhizosphere microbiome composition throughout the growing season; however, the strength of this interaction was minor compared to the effects of location and timepoint (Walters *et al*., 2018). Interestingly, that study revealed a qualitative change in maize microbiomes which occurred in all genotypes between weeks 7 and 8 of development: a bloom of *Pseudomonas*, which increased suddenly in relative abundance from ~3% to over 44%. This bloom was observed only in rhizosphere (not in bulk soil) and persisted for the rest of the growing season. In our study, *Pseudomonas* comprised 2.3% of the rhizosphere community on average, consistent with the results of Walters *et al*. The abundance of *Pseudomonas* varied among genotypes but never averaged more than 3.4% (Figure S10). This suggests that at the time of sampling, no genotype had progressed to the developmental stage that triggers the *Pseudomonas* bloom. Of course, this does not rule out the possibility that more subtle differences in microbiome composition were driven by plant development; however, there is no evidence that phenology was a dominant driver of the observed microbiome differences among genotypes.

Some of the taxa that distinguish hybrids and inbreds (Figure 4) have been linked to positive or negative effects on plant health. For example, *Burkholderia* spp. have been reported to improve growth and stress resilience in diverse plants including maize (Bevivino *et al*., 2000; Mitter *et al*., 2013), and a sizeable minority of maize-associated *Burkholderia* strains appear to be nitrogen fixers (Perin *et al*., 2006). Although maize does not make the root nodules that are central to rhizobial symbiosis in legumes, strains of *Bradyrhizobium* have been shown to increase aboveground biomass in maize by up to 20% (Prévost *et al*., 2012) and can fix nitrogen while free-living (Smercina *et al*., 2019). The yeast-like fungus *Tilletiopsis washingtonensis* produces antifungal metabolites and has shown promise as a biocontrol agent (Urquhart, 1994; Urquhart & Punja, 2002). One ASV identified as *Microdochium sorghi* was strongly enriched in the leaves of hybrids relative to inbreds. This fungus can cause zonate leaf spot disease in maize, although it more commonly infects sorghum (De León, 1978; Korsman *et al*., 2012); however, no symptoms were apparent in the plants in this experiment. Interestingly, a recent study in *Populus* also found an enrichment of foliar fungal pathogens in an interspecific hybrid relative to one of the parent species (Cregger *et al*., 2018).

Without additional data, however, the observed differences in microbiome composition between hybrids and inbreds cannot be reliably translated into differences for microbiome function and/or plant health. This reflects the limitations of the amplicon-sequencing approach. Due to poor taxonomic resolution and a general lack of knowledge about the biology of many microbial taxa, function cannot be reliably inferred from this type of data. Therefore, different approaches are needed to draw conclusions about the importance of microbiome heterosis for community and host function. The true functional consequences of microbiome heterosis would be most clearly assessed by directly manipulating microbiomes and measuring the effects on host phenotype. This approach would also inform us about the direction of causality in the microbiome-heterosis relationship. In this study, we manipulated host genotype and measured the effects on microbial communities; however, given the abundant evidence that plant microbiomes can drastically affect their hosts, the observed effects on the microbiome could feed back and contribute to heterosis of other plant traits. Is microbiome heterosis a meaningless by-product of host hybridization? Or is it a contributing mechanism of hybrid vigor itself? These questions could be addressed using synthetic microbial communities, reinoculation, or soil-conditioning experiments (Ponsford *et al*.; Panke-Buisse *et al*., 2015; Vorholt *et al*., 2017). These important next steps will help to clarify the role of plant-associated microbes in hybrid vigor.

## ACKNOWLEDGEMENTS

We thank G. Marshall, S. Sermons, and J. Torres for assistance with field and lab work. We thank Cathy Herring and the staff at NC Central Crops Research farm for managing the field trials. We thank L. Burghardt and D. Leopold for valuable comments on earlier drafts of the manuscript. All sequencing was done by the North Carolina State University Genomic Sciences Laboratory (Raleigh, NC, USA). M.R.W. was supported by a NSF National Plant Genome Initiative Postdoctoral Research Fellowship in Biology (IOS-1612951) and a NSF EPSCoR RII Track-1 grant (OIA-1656006). This work was funded by a NCSU Plant Soil Microbial Community Consortium grant to J.B.H. and M.R.W.

## AUTHOR CONTRIBUTIONS

MRW, PBK, and JBH planned and designed the research. MRW, JHR, and PBK performed experiments and conducted fieldwork. MRW and JBH analyzed data. MRW wrote the manuscript with edits from PBK and JBH.

## SUPPLEMENTARY INFORMATION

**TABLE S1.** Soil types and crop rotation histories of the two fields at Central Crops Research Station, Clayton, NC, USA. The fields were 1.1 km apart.

| Field | Soil | Rotation (2014-2017) |
| --- | --- | --- |
| A5B | Wagram loamy sand | Squash > Soybean > Corn > Corn |
| G5C | Wagram loamy sand | Corn > Tobacco > Cucumber > Corn |

**TABLE S2.** 49 plant-colonizing ASVs were differentially abundant between the bulk soils of the two fields. The log2FoldChange indicates the direction and magnitude of differential abundance, with positive/negative values describing ASVs that had higher relative abundances in bulk soils of field A5B/G5C, respectively. Column lfcSE notes the standard error of the estimated log2FoldChange. Taxa that could not be identified at the genus level are indicated with parentheses. The “Colonizes” column indicates whether the ASV was observed in leaf samples, rhizosphere samples, or both.

| log2FoldChange | lfcSE | ASV_ID | Kingdom | Genus | Colonizes |
| --- | --- | --- | --- | --- | --- |
| 24.69 | 3.02 | fASV_398 | Fungi | Epicoccum | leaf |
| 23.17 | 3.16 | fASV_148 | Fungi | (Sordariales) | leaf |
| 22.84 | 2.99 | bASV_522 | Bacteria | Ralstonia | leaf |
| 22.79 | 2.89 | bASV_159 | Bacteria | Dyella | both |
| 11.51 | 2.98 | fASV_131 | Fungi | (Sordariales) | both |
| 10.77 | 1.78 | fASV_489 | Fungi | (Cantharellales) | rhizosphere |
| 9.51 | 2.61 | fASV_132 | Fungi | (Sordariales) | leaf |
| 8.51 | 2.31 | fASV_339 | Fungi | Poaceascoma | both |
| 8.41 | 1.74 | fASV_289 | Fungi | Exophiala | leaf |
| 7.93 | 2.04 | bASV_437 | Bacteria | Gaiella | leaf |
| 7.81 | 2.17 | fASV_231 | Fungi | (Mortierellales) | leaf |
| 7.72 | 2.58 | fASV_451 | Fungi | Corynespora | rhizosphere |
| 7.6 | 1.73 | bASV_274 | Bacteria | (Xanthomonadaceae) | rhizosphere |
| 7.42 | 2.1 | fASV_218 | Fungi | Acremonium | both |
| 7.32 | 1.88 | fASV_124 | Fungi | (Sordariomycetes) | rhizosphere |
| 7.24 | 2.97 | fASV_486 | Fungi | Mortierella | both |
| 6.97 | 2.01 | bASV_165 | Bacteria | Rhodanobacter | both |
| 6.96 | 2.55 | bASV_678 | Bacteria | Tumebacillus | rhizosphere |
| 6.79 | 2.07 | fASV_202 | Fungi | (Ascomycota) | both |
| 6.72 | 2.25 | fASV_417 | Fungi | (Tremellales) | both |
| 6.57 | 2 | bASV_737 | Bacteria | Tumebacillus | rhizosphere |
| 6.5 | 2.35 | bASV_610 | Bacteria | Burkholderia | rhizosphere |
| 6.49 | 2.46 | fASV_384 | Fungi | (Fungi) | rhizosphere |
| 6.48 | 2.17 | bASV_163 | Bacteria | Rhodanobacter | leaf |
| 6.36 | 1.71 | bASV_446 | Bacteria | Gaiella | rhizosphere |
| 6.22 | 1.99 | bASV_553 | Bacteria | Massilia | leaf |
| 6.2 | 2.46 | fASV_290 | Fungi | Exophiala | rhizosphere |
| 5.99 | 1.76 | fASV_167 | Fungi | Chloridium | both |
| 5.92 | 1.99 | bASV_442 | Bacteria | Gaiella | both |
| 5.78 | 2 | bASV_571 | Bacteria | Kineosporia | rhizosphere |
| 4.53 | 1.53 | bASV_103 | Bacteria | (Acetobacteraceae) | rhizosphere |
| 4.15 | 1.46 | fASV_106 | Fungi | Microdochium | both |
| 3.78 | 1.3 | bASV_329 | Bacteria | (Chitinophagaceae) | rhizosphere |
| -6.75 | 2.07 | fASV_134 | Fungi | Clonostachys | both |
| -6.87 | 2.38 | bASV_594 | Bacteria | Ramlibacter | both |
| -6.93 | 2.37 | bASV_105 | Bacteria | Rhodoplanes | leaf |
| -7.08 | 2.76 | fASV_150 | Fungi | (Fungi) | both |
| -7.44 | 2.98 | fASV_126 | Fungi | (Sordariomycetes) | leaf |
| -7.51 | 1.92 | bASV_355 | Bacteria | Geobacter | both |
| -8.27 | 2.76 | fASV_498 | Fungi | (GS11) | both |
| -9.17 | 3.11 | fASV_112 | Fungi | (Sordariales) | rhizosphere |
| -9.68 | 2.45 | fASV_160 | Fungi | Chaetomium | both |
| -13.21 | 2.19 | fASV_114 | Fungi | Humicola | rhizosphere |
| -21.72 | 3.13 | fASV_21 | Fungi | Rhizophydium | both |
| -23.11 | 3.14 | fASV_68 | Fungi | Curvularia | rhizosphere |
| -23.48 | 3.13 | fASV_141 | Fungi | Cercophora | leaf |
| -23.98 | 3.14 | fASV_79 | Fungi | Drechslera | rhizosphere |
| -24.14 | 2.79 | fASV_256 | Fungi | Pseudeurotium | rhizosphere |

**TABLE S3.**
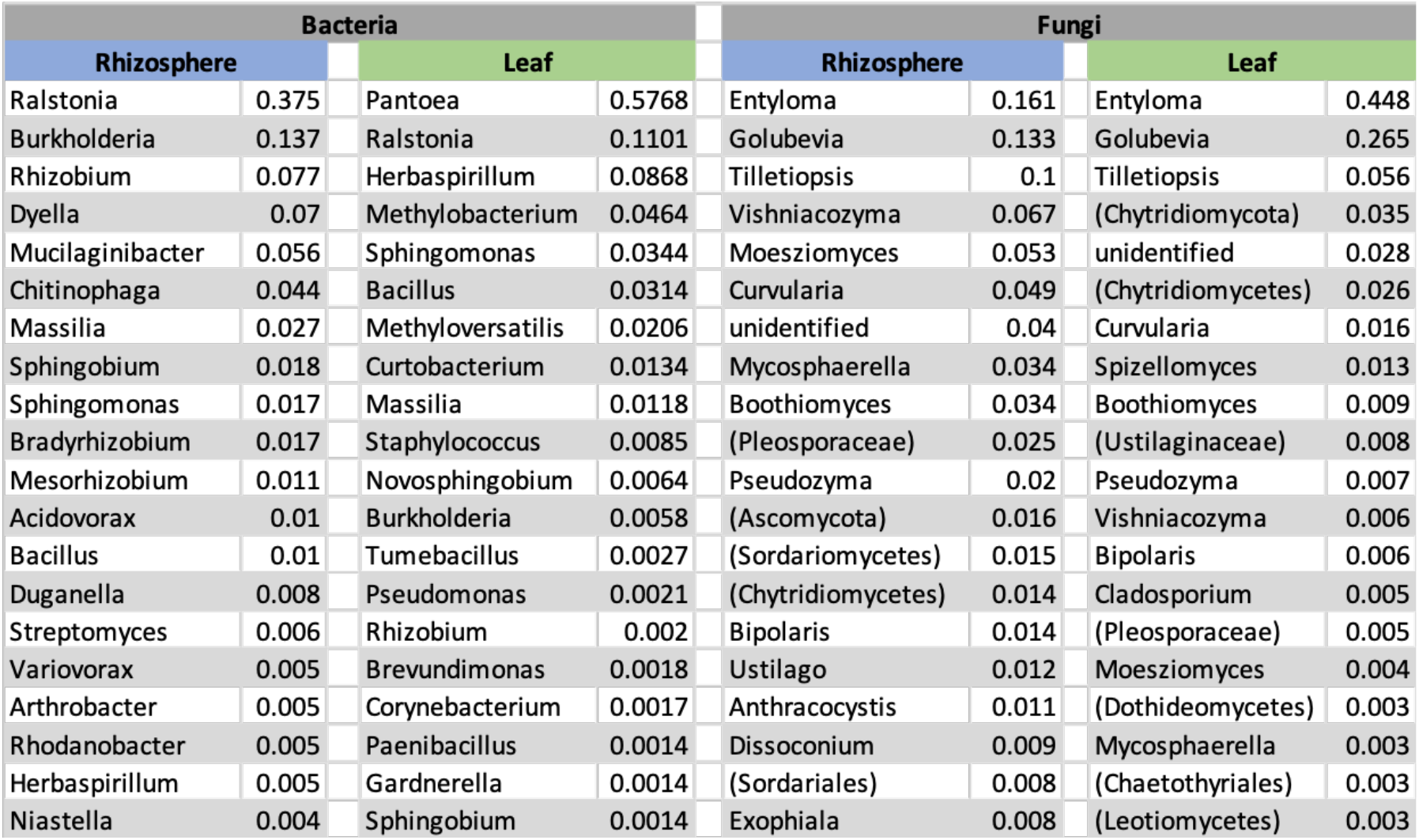
Top 20 genera by relative abundance in leaf and rhizosphere samples. Fungal groups that could not be identified at the genus level are indicated with parentheses.

**TABLE S4.**
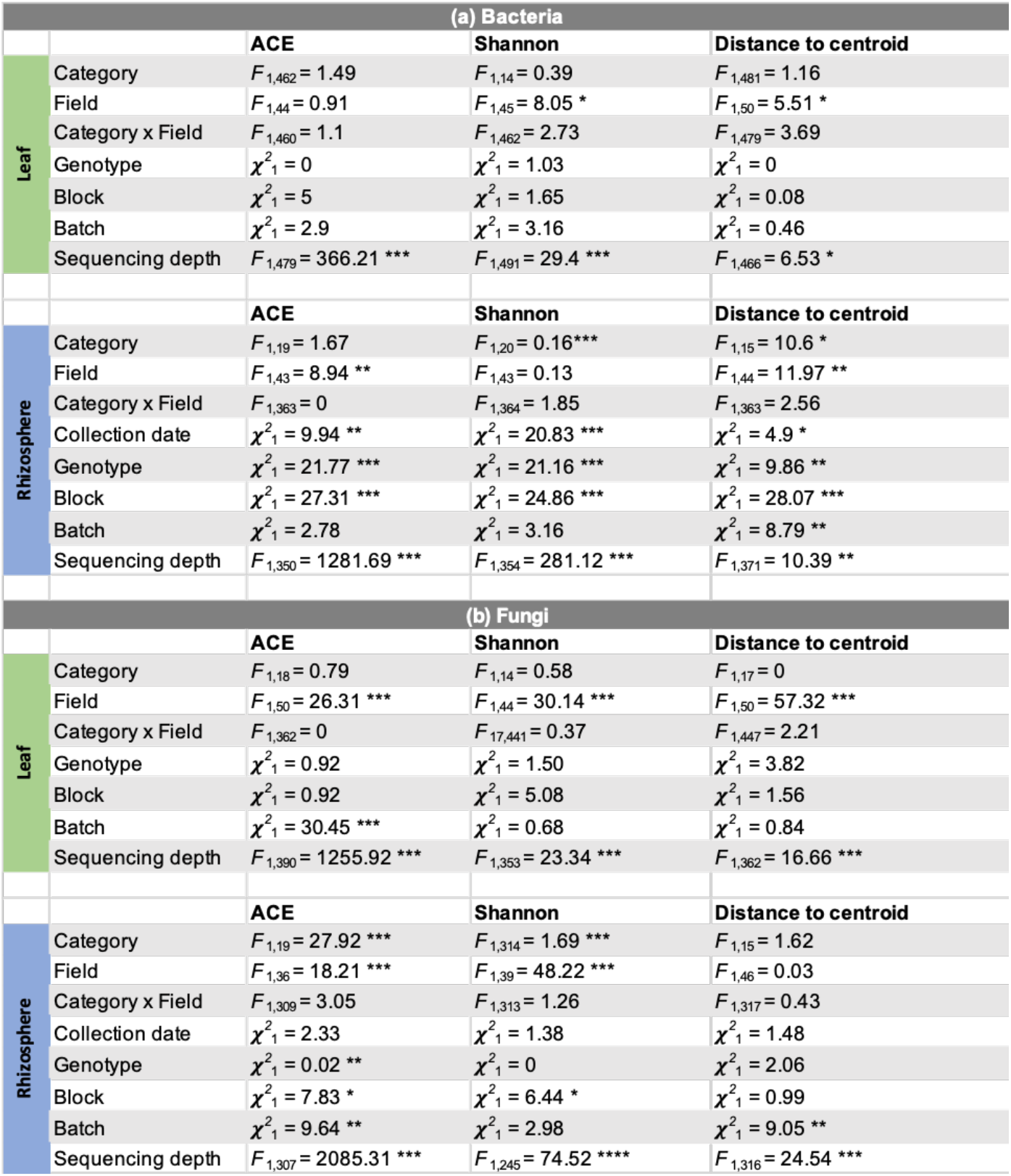
Host genotype influences alpha and beta diversity in maize leaf and rhizosphere microbiomes. ACE (log-transformed) and Shannon metrics were used to quantify community richness and evenness, respectively; Distance to centroid was used to quantify beta diversity (i.e., among-individual variation within genotypes). Bacterial **(a)** and fungal **(b)** communities were tested separately using univariate models and analysis of variance with Type III sums of squares and Satterthwaite’s approximate denominator d.f. Likelihood ratio tests were used for significance testing of random-intercept effects. For rhizosphere fungal communities, one genotype (Oh43) was excluded from the analysis because no individuals survived in one field. *** *P* < 0.001; ** *P* < 0.01; * *P* < 0.05. All *P*-values were adjusted to correct for multiple comparisons using the Benjamini-Hochberg false discovery rate.

**TABLE S5 | ASVs showing significant midparent or “better parent” heterosis in at least one cross.** This table can be downloaded as a separate tab-delimited (.txt) file. Each row summarizes one observation of midparent heterosis (MPH) or “better parent” heterosis (BPH) for one ASV in one hybrid. Column descriptions: Taxon = unique ID of the ASV; Maternal = genotype of maternal inbred line; Paternal = genotype of paternal inbred line; Hybrid = hybrid genotype; MatMean, PatMean, HybMean = mean ASV abundance in the maternal, paternal, and hybrid genotype (residuals of variance-stabilized counts after controlling for Field, Block, Batch, and sequencing depth); MatSE, PatSE, HybSE = standard errors associated with the previously described means; Midparent = expected midparent value of ASV abundance (average of MatMean and PatMean); BetterParent = defined as whichever of MatMean and PatMean is closer to the HybMean; Type = whether heterosis was observed in leaf or rhizosphere; MPH.t.stat, MPH.t.df, MPH.p, MPH.padj = test statistic, degrees of freedom, raw *P*-value, and FDR-corrected *P*-value for T-test comparing HybMean to Midparent; BPH.t.stat, BPH.t.df, BPH.p, BPH.padj = same statistics for T-test comparing HybMean to BetterParent; HetType = BPH if T-test provided evidence for “better parent” heterosis, MPH if T-test supported midparent heterosis but not “better parent” heterosis (ASVs with evidence for neither are not included in this table); Kingdom, Phylum, Class, Order, Family, Genus, Sequence = taxonomic assignment and observed ITS1 or 16S-v4 sequence of the ASV; Colonized = whether the ASV was present in the leaf dataset, rhizosphere dataset, or both.

**TABLE S6 | ASVs with strong loadings on axes of variation with high frequency of heterosis.** This table can be downloaded as a separate tab-delimited (.txt) file.

**TABLE S7.** Permutational MANOVA of fungal and bacterial community composition among genotypes within hybrid/inbred categories. *P*-values were generated from 999 permutations of Bray-Curtis dissimilarities among samples, which were calculated based on the matrix of variance-stabilized counts of amplicon sequence variants (ASVs). Permutations were constrained within Block and within Category.

|  |  | Bacteria |  |  | Fungi |  |  |
| --- | --- | --- | --- | --- | --- | --- | --- |
|  |  | R<sup>2</sup> | pseudo-F test | P | R<sup>2</sup> | pseudo-F test | P |
| Leaf | Sequencing depth | 0.014 | F<sub>1,499</sub> = 7.06 | 0.001 | 0.027 | F<sub>1,478</sub> = 14.73 | 0.001 |
|  | Genotype | 0.043 | F<sub>17,483</sub> = 1.33 | 0.001 | 0.036 | F<sub>17,462</sub> = 1.15 | 0.092 |
|  | Field | 0.021 | F<sub>1,499</sub> = 10.94 | 0.527 | 0.075 | F<sub>1,478</sub> = 40.15 | 0.691 |
|  | Genotype x Field | 0.034 | F<sub>17,483</sub> = 1.03 | 0.166 | 0.036 | F<sub>17,462</sub> = 1.15 | 0.011 |
| Rhizosphere | Sequencing depth | 0.044 | F<sub>1,398</sub> = 24.67 | 0.001 | 0.038 | F<sub>1,329</sub> = 19.41 | 0.001 |
|  | Collection date | 0.082 | F<sub>3,396</sub> = 15.42 | 0.001 | 0.074 | F<sub>3,327</sub> = 12.67 | 0.001 |
|  | Genotype | 0.066 | F<sub>17,382</sub> = 2.17 | 0.001 | 0.063 | F<sub>17,313</sub> = 1.92 | 0.001 |
|  | Field | 0.132 | F<sub>1,398</sub> = 74.37 | 0.001 | 0.226 | F<sub>1,329</sub> = 116.40 | 0.001 |
| | Genotype x Field | 0.033 | $F_{17,382} = 1.08$ | 0.015 | 0.033 | $F_{16,314} = 1.05$ | 0.240 |

**Figure S1.**
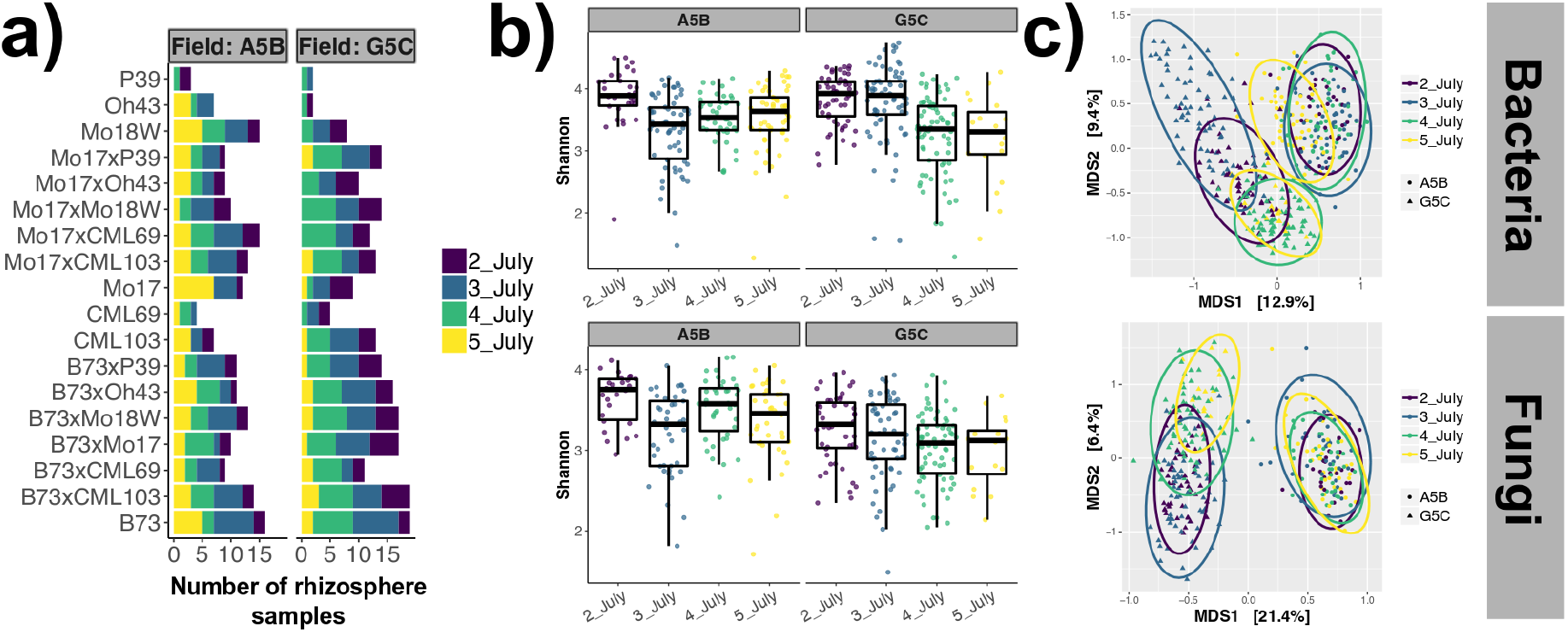
The effect of collection date on rhizosphere microbiome diversity and composition. **(a)** The number of rhizosphere samples in the 16S dataset are shown for each host genotype in each field. Complete replicates were collected on each day so that collection date would not be confounded with either field or plant genotype; however, some genotypes with poor survival are not represented on all dates. **(b)** Shannon diversity varied slightly among collection dates for both bacteria (top) and fungi (bottom). **(c)** Principal coordinates analysis of Bray-Curtis dissimilarities shows that both bacterial and fungal communities shifted in community structure between July 3rd and July 4th, but only in one of the two fields (field G5C).

**Figure S2.**
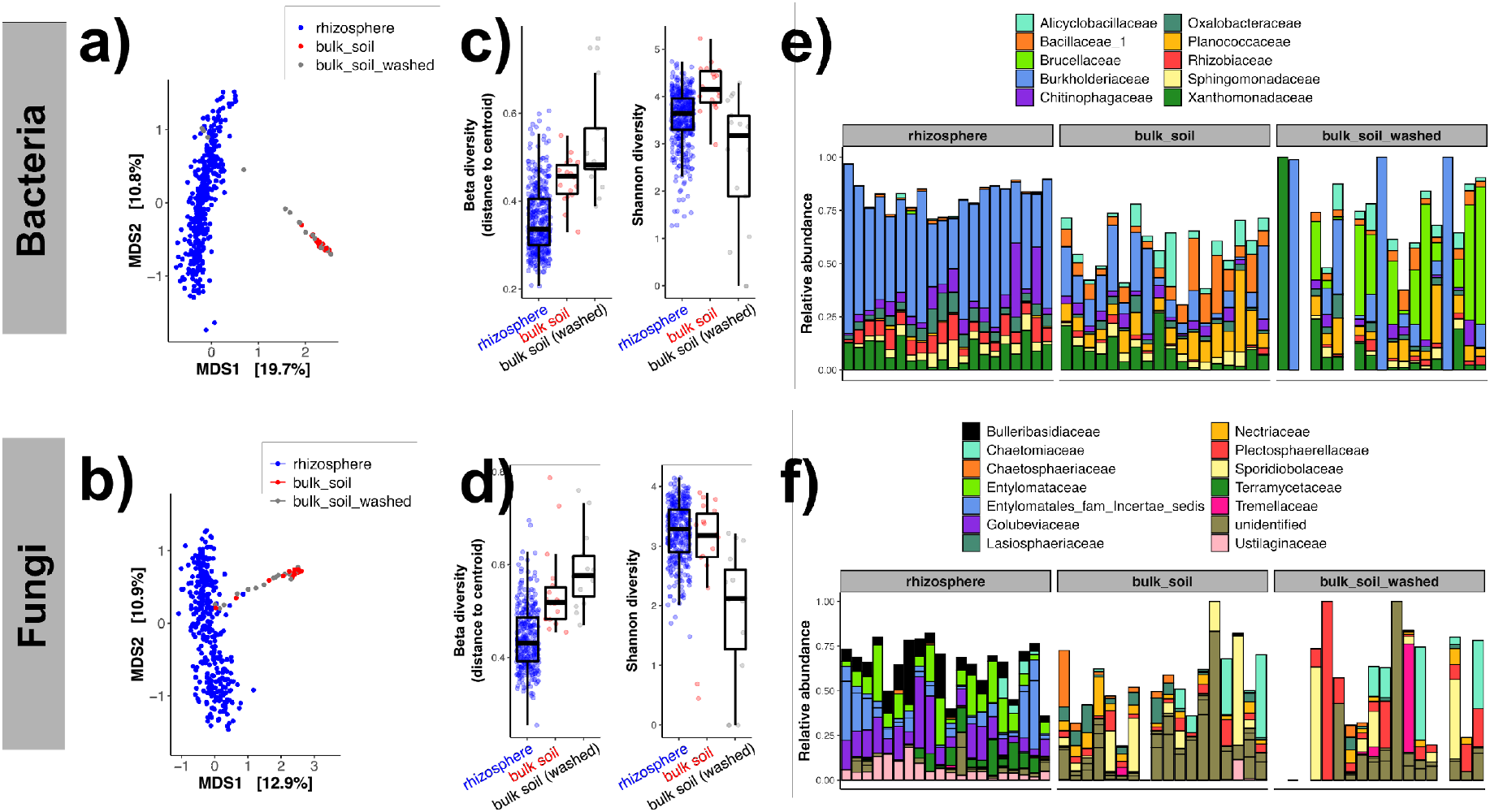
Microbiome characteristics of rhizosphere, bulk soil, and “washed” bulk soil samples. Rhizosphere soil was collected from maize roots using a series of washes and centrifuge steps (see Methods). To assess the effects of this protocol on microbiome readouts, we divided each of 20 bulk soil samples into two aliquots and applied the same protocol to one aliquot prior to DNA extraction. **(a-b)** Multidimensional scaling reveals that bulk soil microbiomes differ substantially in composition from rhizosphere microbiomes, regardless of whether the washing protocol was applied. **(c-d)** For both bacterial and fungal microbiomes, the washing protocol increased beta diversity (left); therefore, the beta diversity values reported for rhizosphere samples in our study is likely an overestimate. The washing protocol decreased alpha diversity (right; measured using the Shannon diversity metric) of both bacterial and fungal communities in bulk soil; therefore, alpha diversity may have been underestimated for the rhizosphere samples in our study if this was due to exclusion or reduction of certain organisms by the protocol. Alternatively, the decrease in alpha diversity may reflect the removal of relic DNA that was artificially inflating diversity estimates, in which case the washing protocol may have increased data accuracy. **(e-f)** Microbiome composition at the family level is shown for randomly selected rhizosphere, bulk soil, and washed bulk soil samples. The 10 most abundant bacterial families and 14 most abundant fungal families are shown. The washing protocol caused notable increases in the relative abundance of Brucellaceae and Plectosphaerellaceae in bulk soil; however, neither of these groups was prominent in the rhizosphere samples.

**Figure S3.**
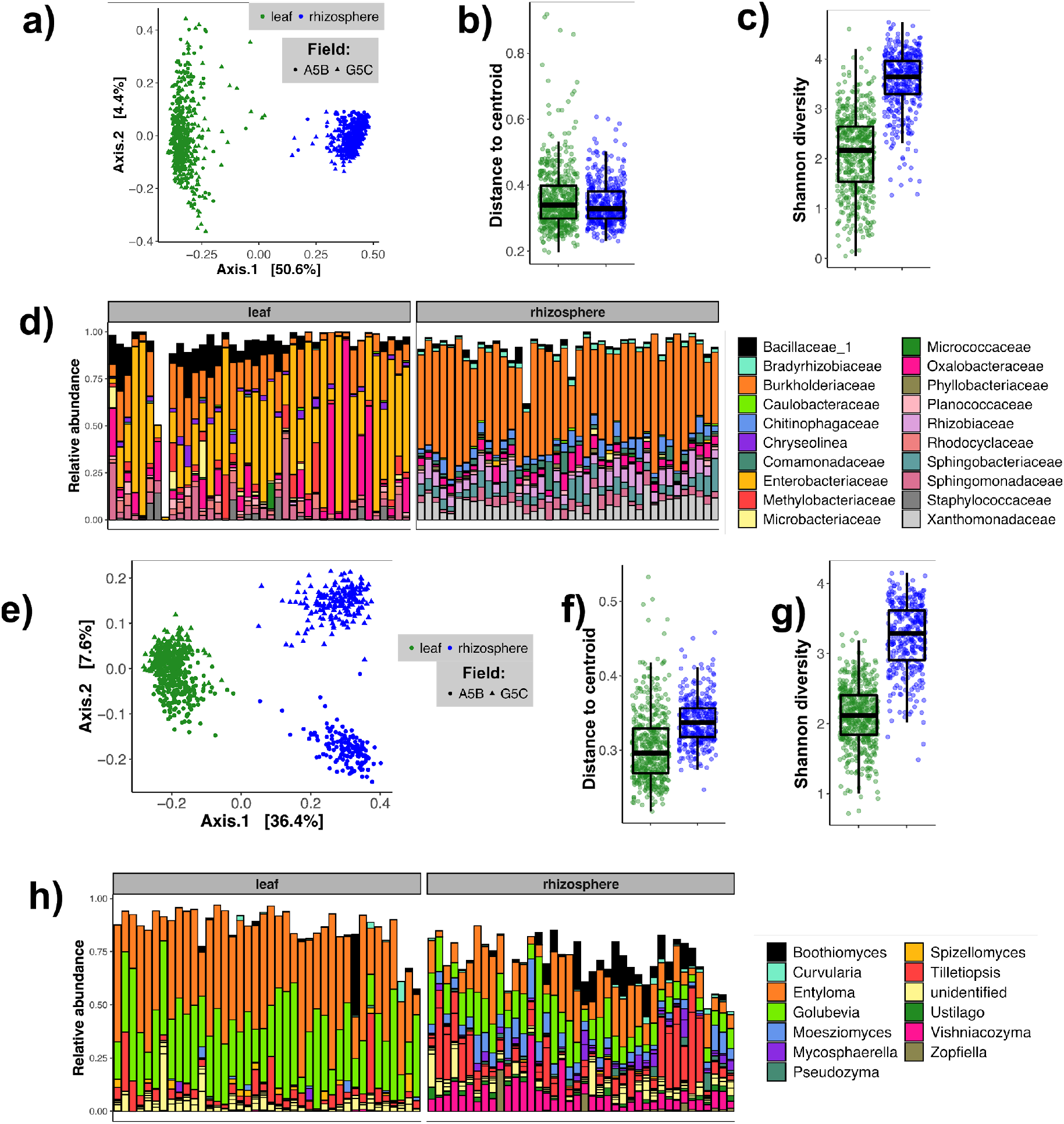
Maize leaves and rhizosphere harbored distinct bacterial (top) and fungal (bottom) microbiomes. **(a,e)** Principal coordinates analysis of Bray-Curtis dissimilarities highlights the difference between leaf and rhizosphere microbial communities. **(b,f)** Beta diversity, measured as distance to centroid (based on Bray-Curtis dissimilarity), was higher in the rhizosphere for fungal communities only. The 75th and 25th percentiles are marked by the top and bottom of each box; the median is marked by the center line; and the length of each whisker is 1.5 times the interquartile range. **(c,g)** Alpha diversity was substantially higher for rhizosphere microbial communities than for leaf communities. Boxplot statistics are the same as for panels (cd). **(d,h)** Leaf and rhizosphere communities were easily distinguished by bacterial and fungal community composition. Each column shows the taxonomic composition of one individual plant, randomly chosen from the dataset.

**Figure S4.**
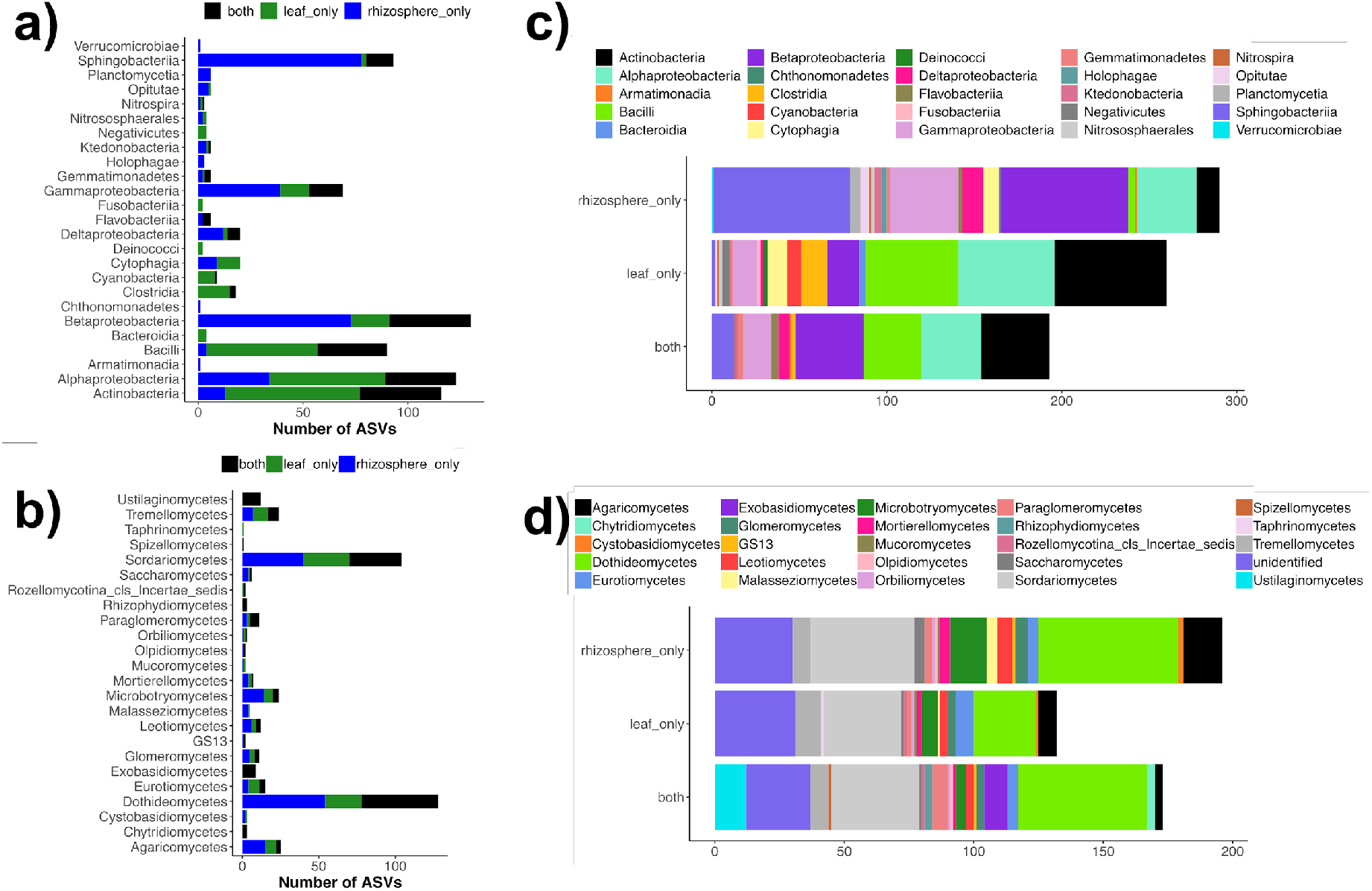
A large minority of bacterial and fungal ASVs were observed in both leaves and rhizosphere. (a-b) For each bacterial and fungal class, the total number of ASVs is shown as well as the number of ASVs that were observed only in leaves, only in rhizosphere, or in both compartments. (c-d) The same data are shown to emphasize the taxonomic makeup of rhizosphere-specific, leaf-specific, and shared ASVs.

**Figure S5.**
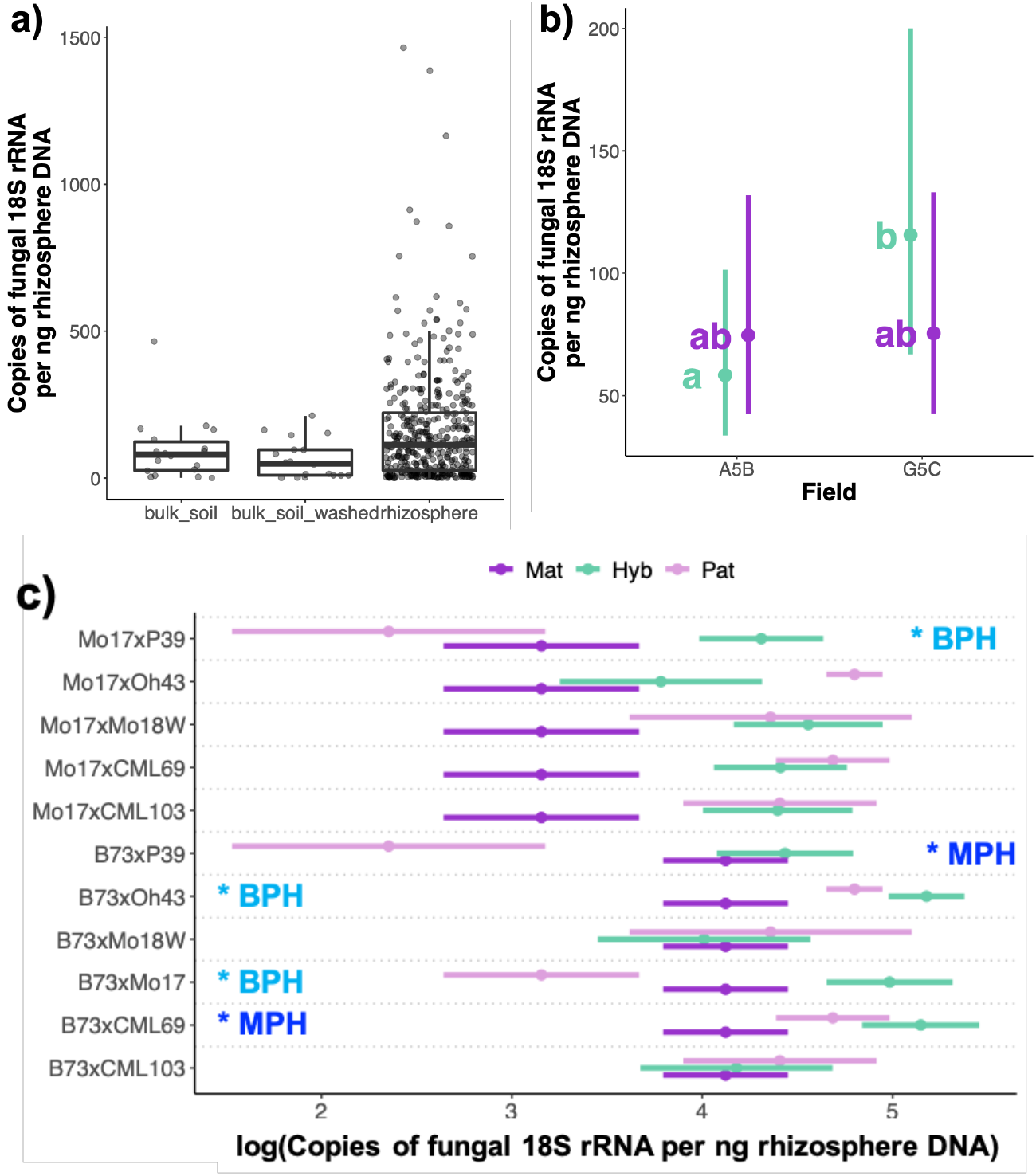
Droplet digital PCR was used to quantify fungal load as the number of copies of fungal 18S rRNA per ng of DNA extracted from soil or rhizosphere samples. **(a)** Fungi made up a greater proportion of the rhizosphere microbiome than the bulk soil microbiome (ANOVA based on linear mixed-effects model of log-transformed fungal load, with Batch as a random-intercept term; *F*_2,425_ = 6.02; *P* = 0.0026). **(b)** For hybrid maize (green) but not inbred maize (purple), fungi made up a greater proportion of the rhizosphere microbiome in field G5C than in A5B (ANOVA, Category x Field *P* < 0.05; post-hoc Tukey’s Honest Significant Differences are shown). However, on average there was no difference between hybrids and inbreds. **(c)** We detected heterosis of fungal load (log-transformed) in five hybrids. Mean values (+/- 1 s.e.m.) are shown for the maternal (“Mat”) and paternal (“Pat”) genotypes and the F1 hybrid (“Hyb”) for each cross. *T*-tests were used to test for midparent and “better parent” heterosis as described in the Methods section of the main text. Statistical support (at *P* < 0.05) for midparent or “better parent” heterosis is noted with “*MPH” and “*BPH”, respectively.

**Figure S6.**
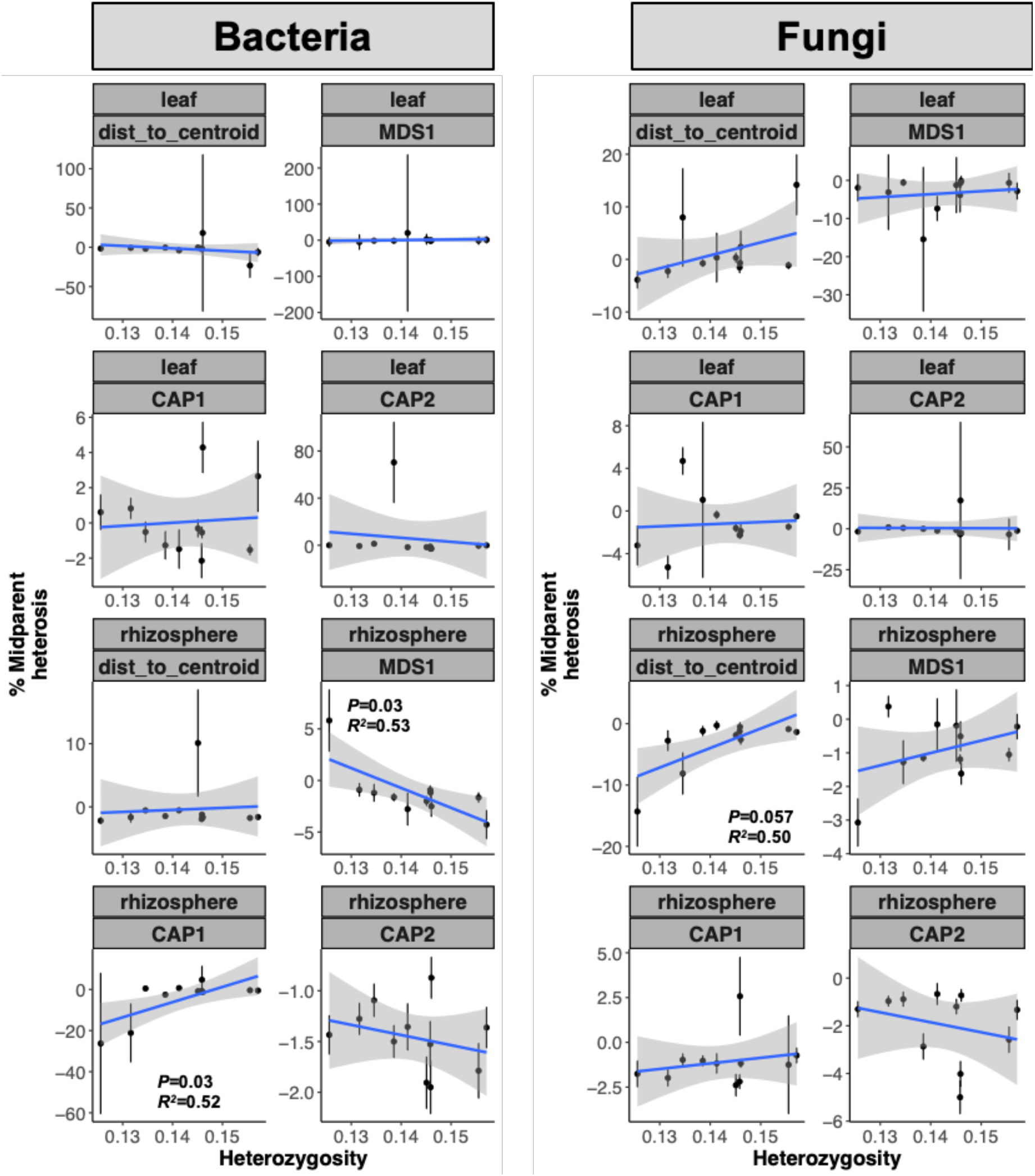
Heterozygosity did not reliably predict heterosis of microbiome composition. For each hybrid, we used genome-wide heterozygosity to predict heterosis of microbiome properties (calculated as the percent change relative to the expected midparent value). Results of linear models are provided; unless otherwise shown, *P* > 0.1 after correction for false discovery rate.

**Figure S7.**
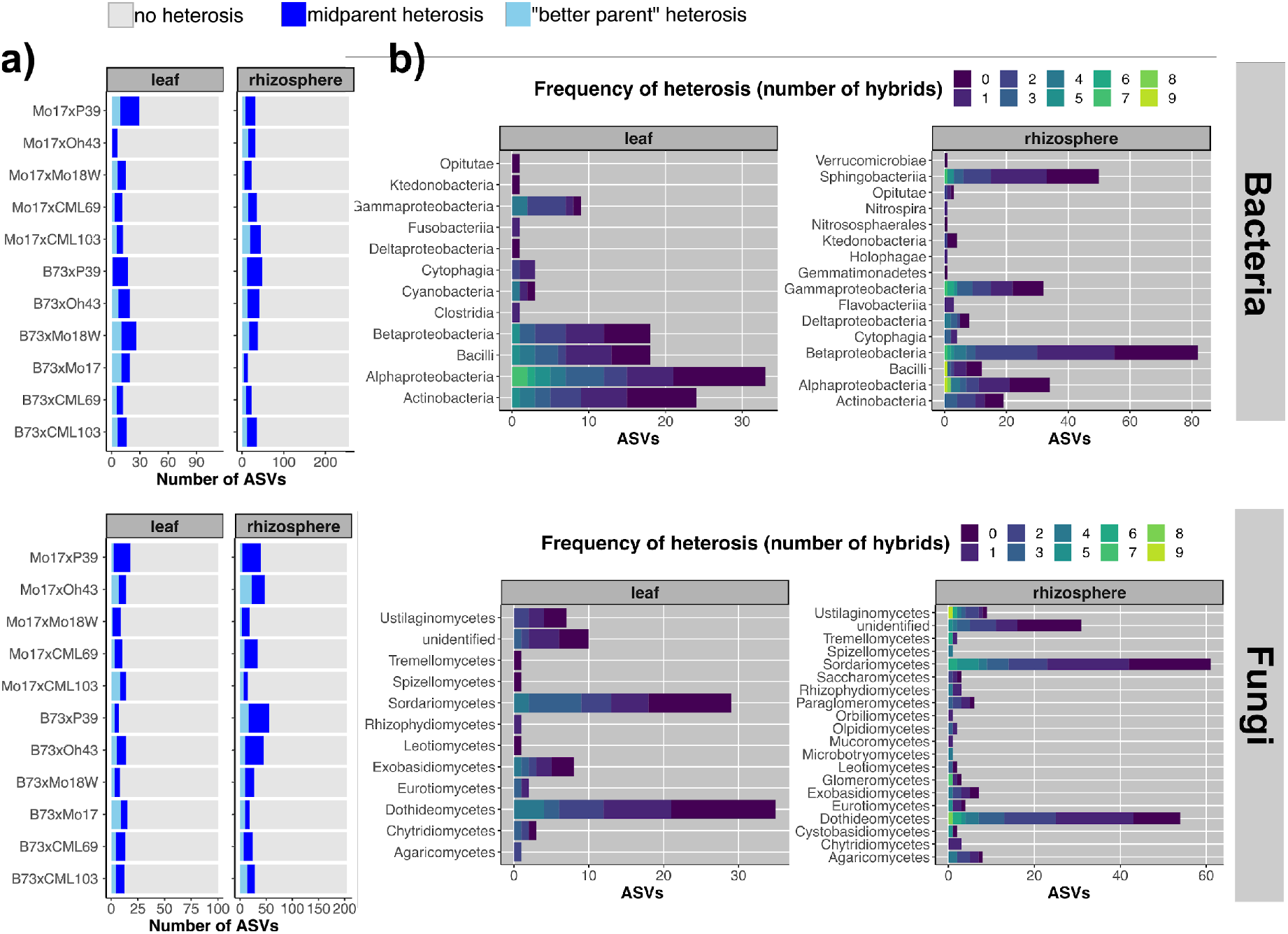
The relative abundances of approximately 35-40% of all rhizosphere ASVs and 17-20% of all leaf ASVs were heterotic in at least one cross. **(a)** The majority of ASVs tested did not show any signal of heterosis in most crosses. ASV heterosis was more common in some hybrids than in others. **(b)** A diverse set of ASVs showed heterosis of relative abundance in at least one hybrid. The *x-*axis portrays the number of ASVs in each taxonomic class, colored according to the number of hybrids in which its abundance deviated from the midparent expectation (Figure 1). Bright colors represent ASVs that showed heterosis in many hybrids; dark colors represent ASVs that were rarely heterotic.

**Figure S8.**
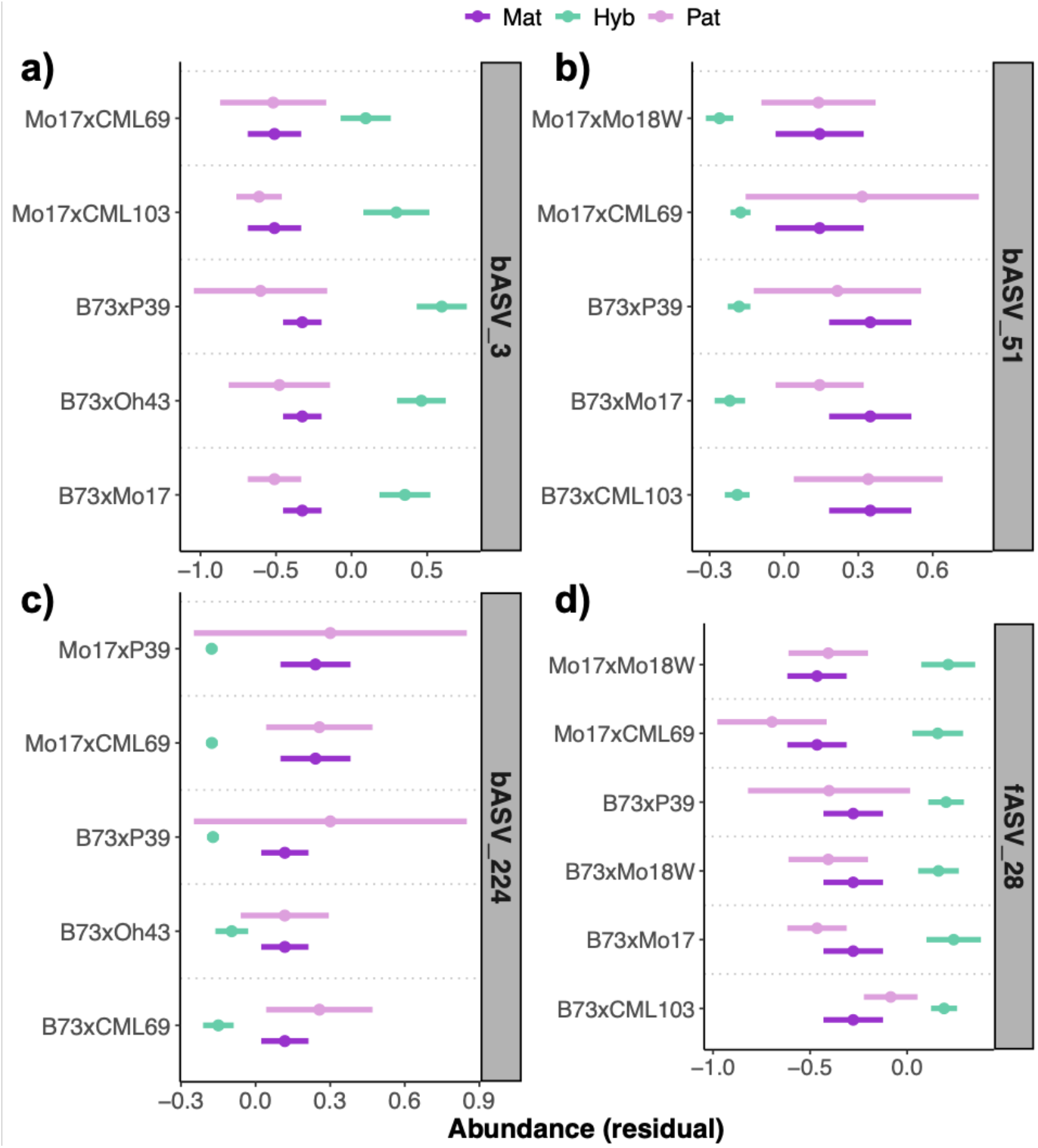
The four taxa showing “better parent” heterosis most frequently. The mean abundance +/- 1 s.e.m. (residuals of variance-stabilized abundances, after controlling for Field, Block, Batch, and Sequencing depth) is plotted for each taxon in the maternal inbred parent (“Mat”), the paternal inbred parent (“Pat”), and the hybrid (“Hyb”). All contrasts between the hybrid and the inbred parents were statistically significant after correction for false discovery rate (*T*-test, FDR < 0.05). **(a)** “bASV_3” belonged to the genus *Pseudonocardia* and showed “better parent” heterosis in the rhizospheres of 5 hybrids. **(b)** “bASV_51” belonged to the genus *Acinetobacter* and showed “better parent” heterosis in the leaves of 5 hybrids. **(c)** “bASV_224” belonged to the genus *Sphingomonas* and showed “better parent” heterosis in the rhizospheres of 5 hybrids. **(d)** “fASV_28” belonged to *Moesziomyces aphidis* and showed “better parent” heterosis in the rhizospheres of 6 hybrids.

**Figure S9.**
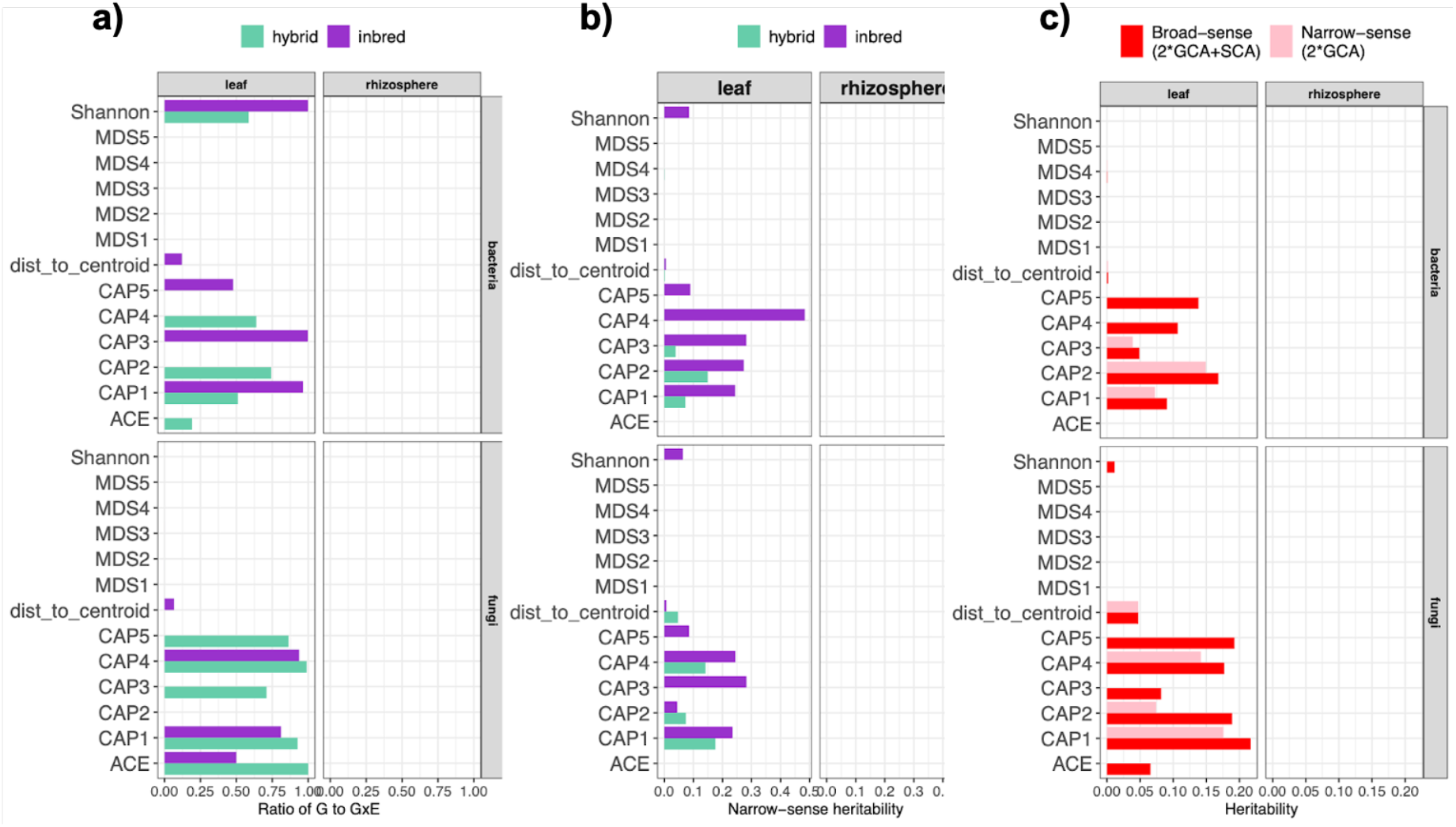
Results of hybrid diallel analysis using linear mixed-effects models for inbreds and hybrids. **(a)** Variance components for Genotype and Genotype-Environment interactions (GxE) were calculated for each trait in hybrids and inbreds, separately. The ratio of G to GxE is plotted for each trait. Higher values indicate lower sensitivity to environmental variation. **(b)** Narrow-sense heritability (due to additive genetic effects) of each trait was calculated for inbreds and for hybrids. **(c)** For hybrids, narrow-sense heritability and broad-sense heritability (non-additive effects) are plotted for each trait. Substantially higher levels of broad-sense heritability indicate an influence of specific parental combinations, beyond the general combining ability (additive effects) of those parents.

**Figure S10.**
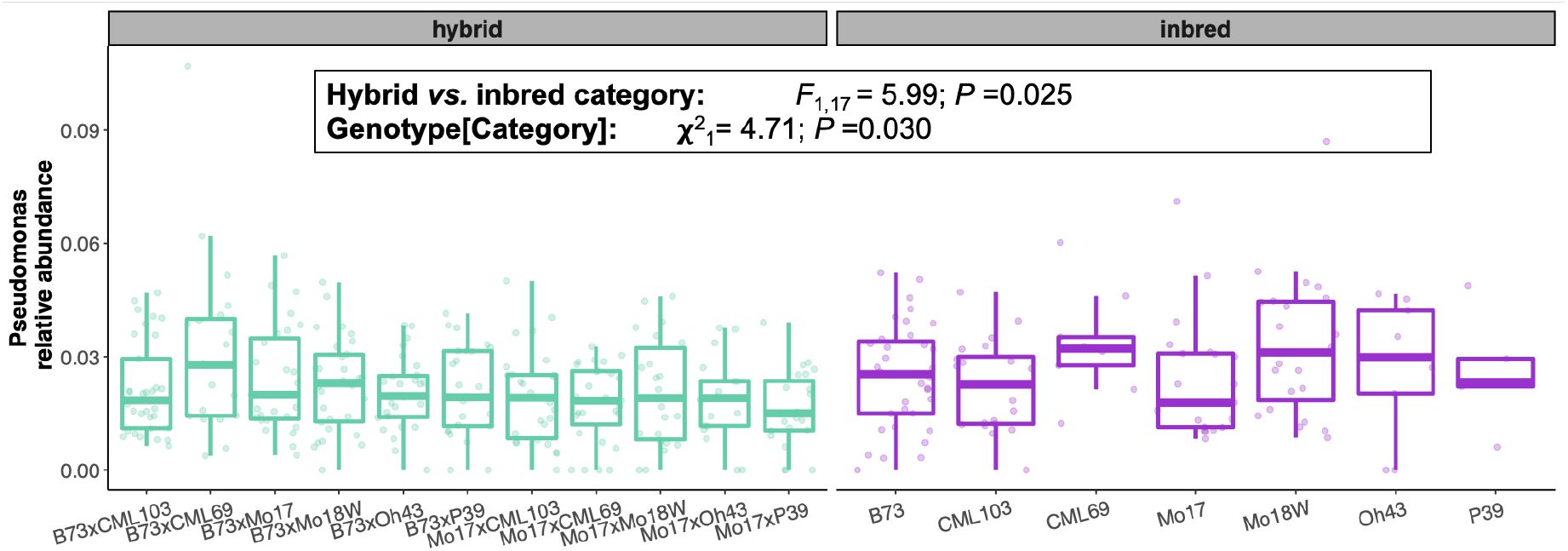
Relative abundance of *Pseudomonas* spp. in the maize rhizosphere 7 weeks after planting. Relative abundances were modeled using hybrid/inbred Category, Field, and their interaction as fixed effect predictors, with Genotype, Block, and Batch as random-intercept terms.

